# Dictys: dynamic gene regulatory network dissects developmental continuum with single-cell multi-omics

**DOI:** 10.1101/2022.09.14.508036

**Authors:** Lingfei Wang, Nikolaos Trasanidis, Ting Wu, Guanlan Dong, Michael Hu, Daniel E. Bauer, Luca Pinello

## Abstract

Gene regulatory networks (GRNs) are key determinants of cell function and identity and are dynamically rewired during development and disease. Despite decades of advancement, challenges remain in GRN inference: dynamic rewiring, causal inference, feedback-loop modeling, and context specificity. To address them, we develop Dictys, a dynamic GRN inference and analysis method which leverages multi-omic single-cell assays of chromatin accessibility and gene expression, context specific transcription factor (TF) footprinting, stochastic process network, and efficient probabilistic modeling of scRNA-seq read counts. Dictys improves GRN reconstruction accuracy and reproducibility and enables the inference and comparative analysis of context specific and dynamic GRNs across developmental contexts. Dictys’ network analyses recover unique insights in human blood and mouse skin development with cell-type specific and dynamic GRNs. Its dynamic network visualizations enable time-resolved discovery and investigation of developmental driver TFs and their regulated targets. Dictys is available as a free, open source, and user-friendly Python package.

## Introduction

Biological molecular networks comprise a compendium of context dependent physical interactions, chemical reactions, and other causal dependencies between mRNAs, proteins, DNA regulatory elements, etc^1^. Together, they define cell identity, gene function, and ultimately phenotypic trait^2–5^. In development, highly ordered, dynamically evolving GRNs are shaped by a repertoire of cell-type specific TFs. They establish regulatory programs by modulating the expression of cell-fate defining genes through binding to their proximal (i.e. promoter) and distal (i.e. enhancer) cis-regulatory DNA elements^6^. Consequently, GRNs have been used to model TF activity, prioritize key master regulators, and understand how they dynamically reshape regulatory programs.

In the past decades, many methods have been proposed to infer GRNs from bulk transcriptomic data alone^4,7–9^. These methods face challenges in distinguishing causation from correlation and direct effects from indirect effects. This motivated the incorporation of other data modalities such as natural genetic variations as causal anchors^10,11^ or TF binding (e.g. from ChIP-seq or chromatin accessibility) as mechanistic evidence^12,13^. However, these bulk-based approaches demand considerable cost for sufficient transcriptomic samples and consequently statistical power, severely limiting their practical affordability, let alone comparative studies^14,15^ such as context specific or dynamic GRNs.

Single-cell RNA sequencing (scRNA-seq) has overcome some of these limitations by cost-efficient profiling the transcriptional programs of thousands of cells in parallel and led to new computational developments of GRN inference. SCENIC, one of the first such methods, reconstructed population-level GRNs with random forest regression by integrating scRNA-seq data with conservation-based putative TF binding information in gene promoters^16^. CellOracle further inferred cell-type specific GRNs with linear or bagging regression from scRNA-seq and population-level chromatin accessibility^17^.

However, single-cell GRN inference faces several challenges, some of which are decades-old and unsolved^2,4,5,11,18^ (**Supplementary File 1**). In fact, these approaches are based on steady-state regression models and consequently cannot account for feedback loops^19^ or detection noise^11^, which is conversely aggravated and confounded by single-cell sparsity^20^. In addition, neither SCENIC nor CellOracle accounts for cell-type specific TF binding in distal regulatory elements such as enhancers, or models the (pseudo-)time-resolved dynamic GRN rewiring in continuous processes, such as development or differentiation^21^. Single-cell multi-omic assays can jointly profile gene expression and chromatin accessibility^22–24^, but they are still yet to be utilized for mechanistic insights in GRN inference. Altogether, these challenges hinder an accurate, reproducible, context specific, and time resolved GRN inference.

Here we propose Dictys to address these challenges and to take advantage of recent multi-omics technologies using context specific TF footprinting, stochastic process model, probabilistic programming, and dynamic network. These advances improve GRN inference quality and enable the comparative analyses of context specific GRNs and dynamic GRNs along differentiation paths thanks to dedicated network visualizations. We demonstrate their utility in blood and skin systems as developmental contexts by rediscovering known key regulators and by uncovering and prioritizing novel regulator candidates and their target genes.

## Results

### Overview of Dictys

Dictys is an integrative Python package for network inference, analysis, and visualization that takes in joint or separate profiles of scRNA-seq and bulk or scATAC-seq data to uncover master regulators, their target genes, and their rewiring that are associated with continuous processes such as development. Dictys can infer context specific networks among annotated cell groups or biological contexts (e.g. clusters or sorted populations) and dynamic networks along continuous processes (e.g. pseudo-time, RNA-velocity, or clonal-based trajectories). Dictys offers network-based analyses and visualizations for the discovery and inspection of each TF’s regulatory activity shifts, of which TF expression level is only a proxy and cannot fully capture.

To reconstruct a context specific GRN for each group of cells, Dictys first infers TF binding sites in regulatory regions (i.e. promoters and enhancers) from TF footprints in pseudo-bulk or bulk chromatin accessibility data (**Fig. 1ab, Methods**). TF footprints are much shorter regions compared to chromatin accessibility peaks and can mitigate false positive binding sites^25^. This selection step prioritizes context specific regulatory TF-target gene links as the TF binding network based on inferred binding and proximity.

**Fig 1.**
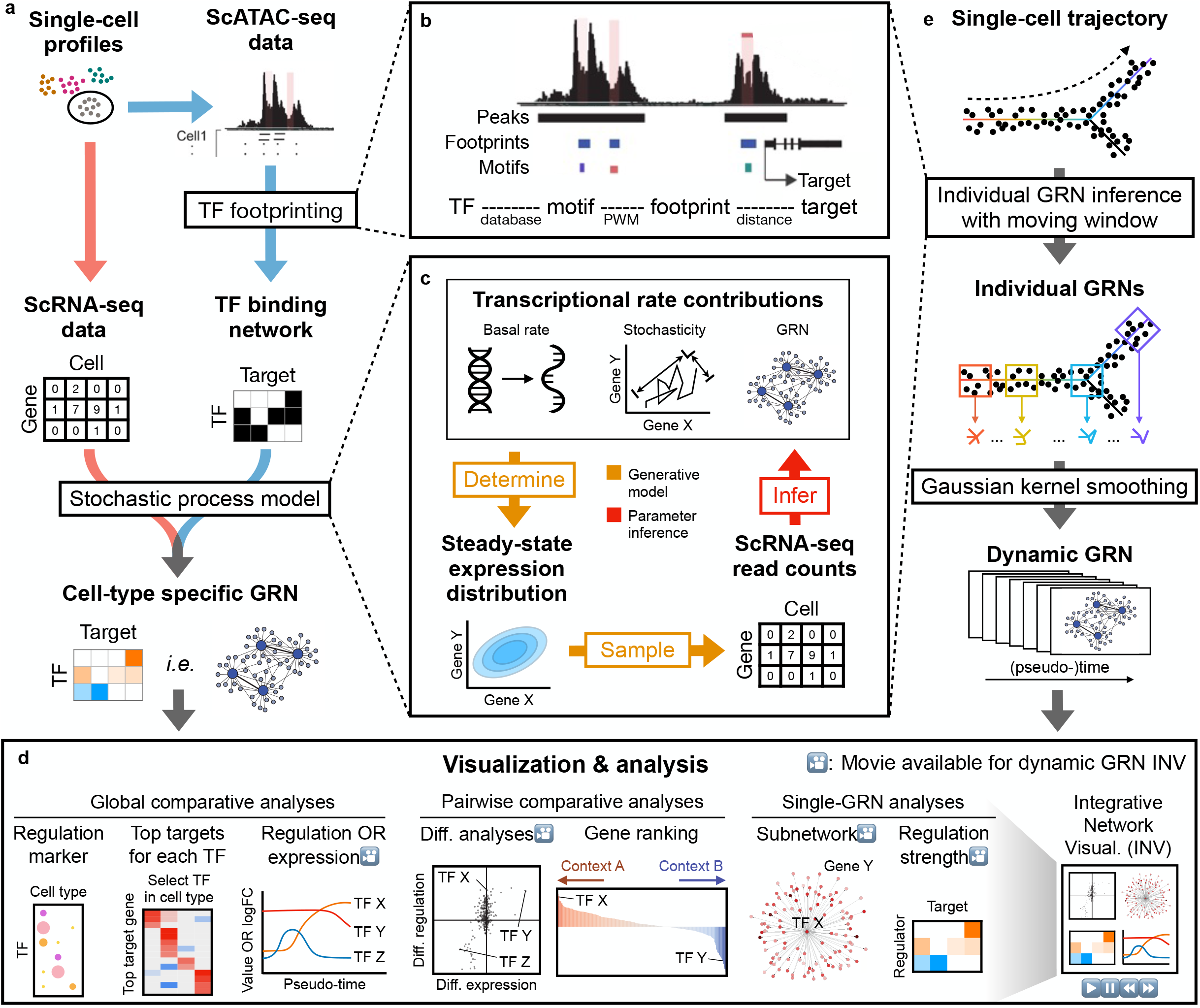
An overview of Dictys to reconstruct and analyze context specific and dynamic GRNs from single-cell transcriptome and chromatin accessibility profiles. **a** Schematic overview of context specific GRN reconstruction for a given cell subpopulation. **b** Initial TF binding network reconstruction from scATAC-seq data with TF footprinting. Each TF was linked to its potential target genes through its motifs and context specific active footprints. **c** Context specific GRN reconstruction from scRNA-seq data with stochastic process models under the TF binding network constraint. The kinetic model parameters including the GRN analytically determine the steady-state distribution of single-cell gene expression. From the steady-state distribution, scRNA-seq sparsely samples the read count matrix with strong technical noise. The full process was characterized in a generative model for parameter inference (including for the GRN) with probabilistic programming. **d** Comparative GRN visualizations and analyses of (left-to-right) multiple (≥ 2), two, or single static GRN(s), and of dynamic GRN in movie format. **e** Schematic illustration of dynamic GRN reconstruction along single-cell trajectories.

Dictys then refines this initial TF binding network using single-cell transcriptome data (**Fig. 1c, Methods, Supplementary File 1**). We model single-cell transcriptional kinetics to allow for feedback loops, using the Ornstein-Uhlenbeck (OU) process^26^ with empirical contributions from basal transcription, direct GRN by TF binding, and stochasticity. Its steady-state distribution^27^ then characterizes the biological variations in single-cell expression. Conversely, single-cell technical variation/noise is modeled with sparse binomial sampling^28^. This proposed generative process is trained on scRNA-seq read counts to infer all kinetic and stochastic parameters, including the GRN, with the probabilistic programming framework Pyro^29^. The resulting GRN is further scale-normalized to account for variance underestimation bias due to single-cell sparsity^20^. Such kinetic GRNs can also simulate every gene’s perturbation as a basal transcriptional rate change, and analytically derive the *total-effect (direct + indirect) GRN* as the consequential change of steady-state expression for other genes.

Dictys includes a suite of functions to understand and compare context specific networks. By identifying the set of target genes for each TF (regulon), the TF *regulatory activity* can be quantified as target gene count based on the recovered network. This is in contrast with gene-level analysis based only on the TF *expression level* defined as Counts-Per-Million (CPM, **Fig. 1d, Methods**). By comparing context specific networks at the global level, Dictys can prioritize *regulation marker* TFs based on context specific overabundance of target count (as opposed to *expression marker* genes from CPM overabundance) and summarize them visually in a dot plot. The regulatory programs of regulation markers are further established and visualized with a heatmap of top target genes in the relevant context. Between two context specific networks, Dictys can unveil different modes of TF activity shift based on recovered *differential regulation* (i.e. logFC in regulatory activity) and *differential expression* (i.e. logFC in CPM). This relationship can be visualized in a scatter plot to uncover unique TFs with strong changes in regulatory activity but not expression, which will be missed based on expression information alone. By further integrating these two differential axes (mean logFC), Dictys provides an *integrative TF ranking* visualized as a bar plot. At the single-network level, Dictys also visualizes individual regulons in a network graph or heatmap format for in-depth investigation.

Dictys infers and analyzes (pseudo-)time-resolved dynamic GRNs to dissect gene regulation variations in continuous processes like development with a single snapshot experiment. Along the provided trajectory, Dictys first defines a moving window to subset cells into overlapping small (∼1000 cells) subpopulations, and then reconstructs a static GRN for each subpopulation and consequently the dynamic GRN with Gaussian kernel smoothing (**Fig. 1e, Methods**). With dynamic GRN, Dictys defines the *regulatory activity curve* for each TF as their regulatory activity variation over time. Dictys then discovers TFs with a highly variable regulatory activity curve in monotonic or transient modes, and performs investigative analysis and animation for individual genes and regulations with integrative network visualization (INV) (**Fig. 1d**).

In total, Dictys provides an inference, visualization, and analysis framework for context specific and dynamic GRNs from single-cell transcriptome and chromatin accessibility profiles that addresses several limitations of existing approaches in context specificity, time resolution, feedback loop, and single-cell detection noise.

### Dictys uncovers cell-type specific GRNs for TF discovery and interrogation in human hematopoiesis

To demonstrate the utility of Dictys in developmental contexts, we obtained a human blood dataset containing scRNA-seq and scATAC-seq data of bone marrow mononuclear cells^30^ (**Fig. 2a, Fig. S1a**). We first identified cell-type specific TF footprints, as shown for three lineage-defining TFs as examples (**Fig. S1b, Methods**). Quality assessment identified sufficient footprints that allowed the reconstruction of cell-type specific GRNs for 12 cell types across human hematopoiesis (**Fig. S1c**).

**Fig 2.**
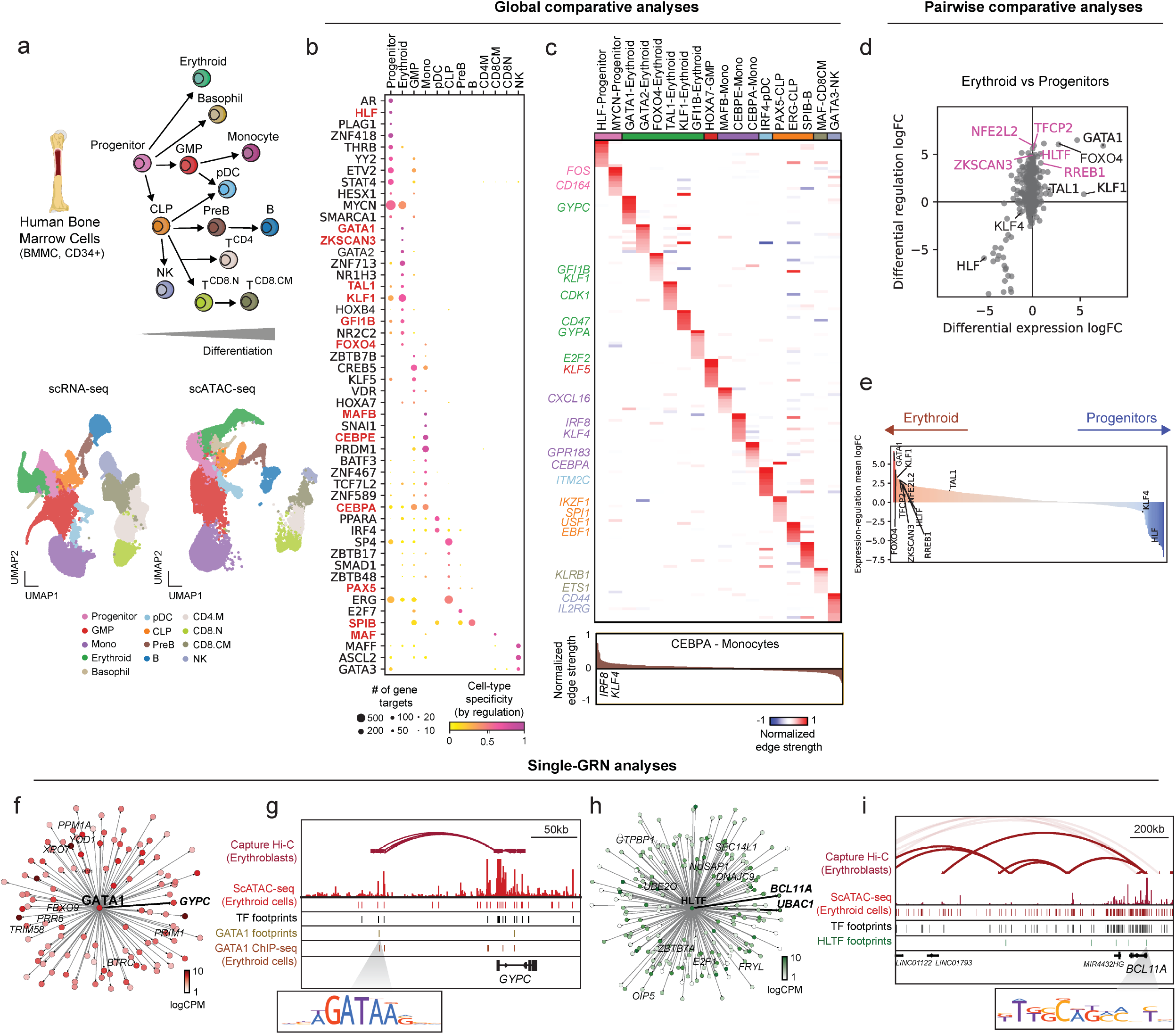
Dictys uncovers cell-type specific GRN, novel and established TFs and regulations in human hematopoiesis. **a** Hierarchical cartoon representation (top) and UMAP embedding (bottom) of the human hematopoietic cell types identified in scRNA-seq and scATAC-seq profiles of Bone Marrow Mononuclear Cell (BMMC) and CD34+ primary samples. **b** Regulation marker discovery based on target count (node size) and its specificity (node color) uncovered known (red) and novel TF markers for each cell type. **c** Top: Heatmap of the regulation strengths from each prominent TF marker (column) to its top 10 activated targets (row) in its corresponding cell type. Target gene names (rows) are colored by cell type. Bottom: regulation strengths from CEBPA to its potential targets in Monocytes as an example. **d** Scatter plot of TF differential expression (X) and differential regulation (Y) logFCs between Erythroid and Progenitor cells. Known TFs with strong activation only in differential regulation are annotated in pink. **e** Integrative TF ranking between Erythroid (red) and Progenitor (blue) cells using the mean logFC of differential expression and differential regulation. **fghi** Subnetwork plots of select TFs’ (GATA1, **f;** HLTF, **h**) inferred targets and IGV snapshots with experimental data of select regulations (GATA1-GYPC, **g;** HLTF-BCL11A, **i**) in Erythroid cells. Subnetwork node colors indicate gene expression. More strongly regulated targets are closer to central node.

By comparing GRNs across all these cell types, Dictys identified regulation marker TFs with clear cell-type specific regulatory activity (i.e. overabundance in target count), including major lineage-defining TFs in Stem/Progenitor (HLF, GATA2)^31^; Erythroid (GATA1, GATA2, TAL1, KLF1)^32–35^; Monocytes (CEBPA, CEBPE, MAFB)^36–38^; and B cells (SPIB, PAX5)^39,40^ (**Fig. 2b, Methods**). Notably, some factors such as CREB5, a key granulocyte regulator in granulocyte-monocyte progenitors (GMPs)^41^, were identifiable only with regulatory activity but not expression (**Fig. S1d**). Moreover, Dictys unveiled the cell-type specific regulatory programs of these regulation marker TFs based on their top activated targets (**Fig. 2c**). This analysis uncovered regulation of surface markers (e.g. GATA1-*GYPC*^42^, KLF1-*CD47*^43^ in Erythroid cells, MYCN-*CD164*^44^ in Progenitors, etc) and others (e.g. GFI1B-*E2F2* in Erythroid cells^33,45^). This demonstrates that cell-type specific GRNs could identify regulation marker TFs and their regulatory programs beyond mean expression-based analyses.

We next contrasted GRNs from early to late stages of hematopoiesis to dissect the regulatory shifts of each TF based on differential regulation (in target count) and differential expression (in CPM). Although both differential analyses of Erythroid versus Progenitor cells recovered many established TFs (e.g. GATA1, GFIB, FOXO4 for Erythroid cells and HLF, KLF4 for Progenitors^31–35,46^), several TFs with Erythroid-specific functions exhibited much stronger logFCs in differential regulation, such as TFCP2 and RREB1 (hemoglobin regulators^47,48^), HLTF, ZKSCAN3, NFE2L2, etc^49–51^ (**Fig. 2d, Supplementary Table 1**). Accordingly, integrative TF ranking with the mean logFC of differential expression and differential regulation could improve ranking quality, especially for known TFs that were weak in one axis (e.g. TFCP2, NFE2L2, and ZKSCAN3, **Fig. 2e, Fig. S1e**). In summary, differential regulation could identify TFs with cell-type specific functions undetectable in differential expression and provide independent information for comparative TF ranking.

For each TF, Dictys could also uncover its cell-type specific regulatory program and mechanistic insights at gene and locus levels. In Erythroid cells, GATA1 was found to regulate erythroid-specific genes, such as the surface marker *GYPC*^42^ and the nuclear export protein XPO7^52^ (**Fig. 2f, Supplementary Table 2, Methods**). This was corroborated by chromatin conformation and GATA1 binding data^53^ overlayed with GATA1 footprints and scATAC-seq fragments around the *GYPC* genomic locus (**Fig. 2g**). HLTF, a known SWI/SNF chromatin remodeler in pro-erythroid cells implicated in genomic instability in acute myeloid leukemia^50,54^, also had elevated regulatory activity in Erythroid cells (**Fig. 2de**) with prominent inferred targets such as *BCL11A* and *UBAC1*^35,55^ (**Fig. 2hi**).

The above analyses were generalizable to other hematopoetic lineages. For example, we (re)discovered known (SPIB, PAX5) and novel (SMAD2) TFs and their putative targets in B cells^39,40^ (**Fig. S1f-i, Supplementary Tables 1-3**). Taken together, Dictys regulatory analyses recovered known and potentially novel insights of hematopoiesis hidden in traditional expression-based analyses.

### Dictys refines GRN inference by leveraging multi-omic data and transcriptome-chromatin accessibility associations in mouse skin

Next, we sought to demonstrate how Dictys could refine GRN inference by leveraging recent joint transcriptome-chromatin accessibility data. To this end, we reanalyzed a SHARE-seq dataset profiling mouse skin development^22^ (**Fig. 3a, Fig. S2a**). Using this multimodal data, we restricted the initial TF binding network to those with population-level correlation between peak chromatin accessibility and target gene expression (**Fig. 1b, Methods**). Dictys identified regulation marker TFs distinct from expression markers^56–63^ as before (**Fig. 3b, Fig. S2b**). As an example, differential regulation analysis between basal epidermis cells and hair follicle TAC-1 revealed TFs that were known but lowly ranked or with opposite effects in differential expression, e.g. Jund, Junb, Fos, and Thrb^64,65^ (**Fig. 3c, Supplementary Table 1**). These TFs were ranked as more cell-type specific by the integrative TF ranking than by differential expression alone (**Fig. 3d, Fig. S2c**). Therefore, Dictys can be used with recent multi-omics data and leverage transcriptome-chromatin accessibility associations to refine GRN inference.

**Fig 3.**
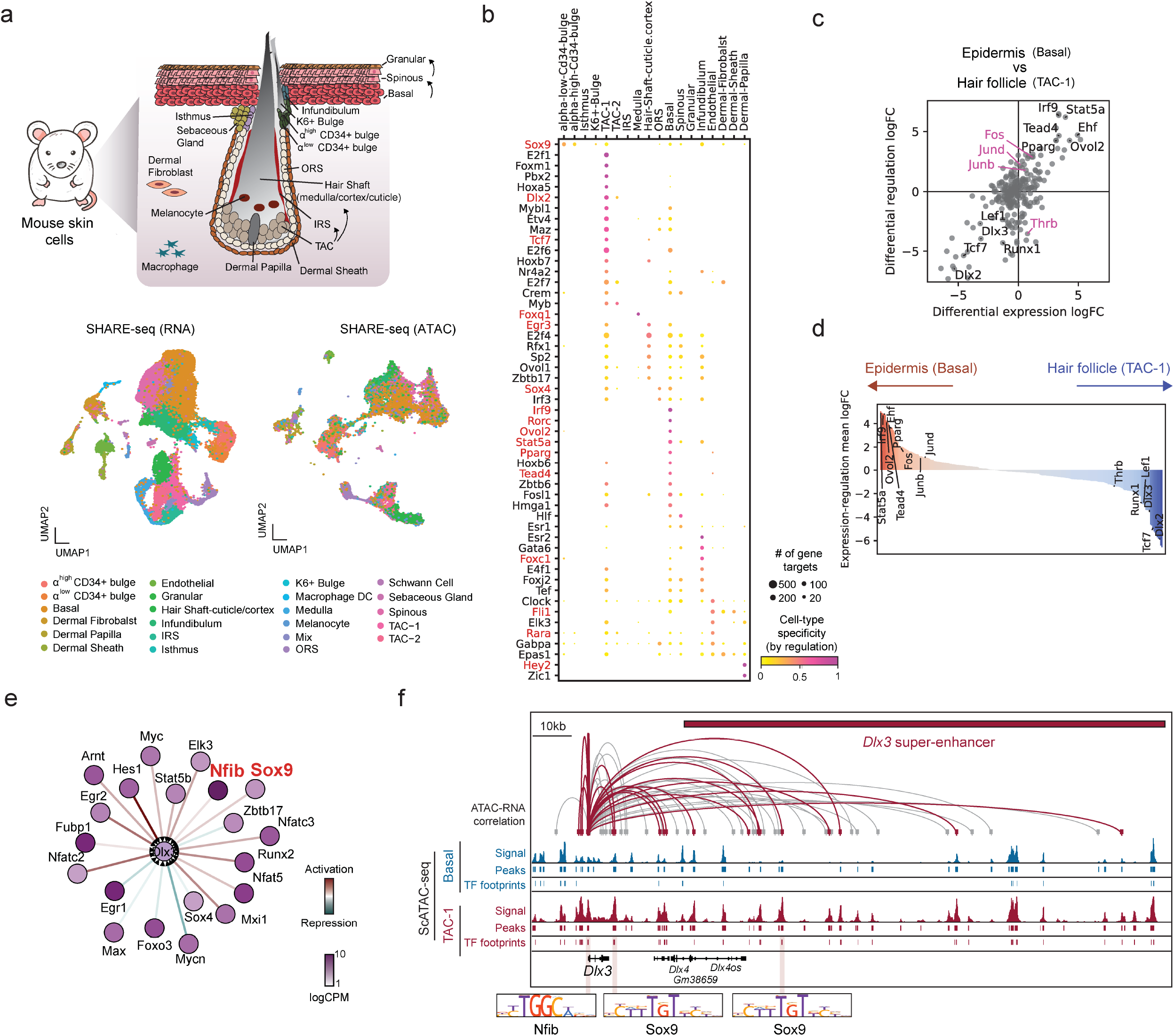
Dictys refines GRN inference with multi-omic data from mouse skin. **a** Top: Cartoon representation of the mouse skin cell types. Bottom: UMAP embedding of cells based on transcriptome and chromatin accessibility profiles. **b** Regulation marker discovery uncovered known (red) and novel TF markers for each cell type based on target count (node size) and its specificity (node color). **c** Scatter plot of TF differential expression and differential regulation logFCs between epidermal (Basal) and hair follicle (TAC-1) cells. Known TFs with strong activation only in differential regulation are annotated in pink. **d** Integrative TF ranking between epidermal (Basal, red) and hair follicle (TAC-1, blue) cells. **e** Subnetwork plot of the inferred regulator TFs of *Dlx3* in TAC-1. Prominent regulators of hair follicle development are annotated in red. **f** IGV snapshot of epigenomic information on *Dlx3* super-enhancer supporting its regulation by TFs in TAC-1 cells. Genomic regions significantly associated with *Dlx3* expression at population level (top arches, P<0.1, height proportional to logP) are annotated based on whether they are accessible in TAC-1 (red), alongside cell-type specific scATAC-seq signal, peaks, and footprints, and prominent TF regulator Nfib and Sox9 motifs.

Transcriptome-chromatin accessibility associations could also help prioritizing and uncovering individual regulations hidden otherwise. However, they do not provide any information on the potential regulators that may bind to these regions, or their regulatory strength in mediating cell-type specific gene expression. Our network analyses based on the recovered regulons provide a solution to this and allow to prioritize peak-to-gene interactions. For example, the previously reported *Dlx3* super-enhancer, present in TAC-1 but not Basal cells, encodes DNA regulatory motifs hinting to potential regulators^22^.

With the TF binding network refinement from transcriptome-chromatin accessibility associations, Dictys identified 14 putative activators and 6 repressors in TAC-1 (**Fig. 3e, Supplementary Table 2**), including activating regulators Sox9^22,66^ and Nfib^67^ that were notably absent if no refinement was applied (**Fig. S2d**). Importantly, Dictys could resolve the epigenomic information underlying these regulatory connections, such as *Dlx3*’s mRNA-associated Sox9 and Nfib footprints in enhancer and promoter areas, respectively, that were absent in Basal cells (**Fig. 3f**). In total, Dictys could refine GRN inference based on new multimodal data and with custom filters for initial TF binding network depending on biological focus.

### Dictys outperforms existing methods in quantitative benchmarks

GRN inference benchmarking remains challenging due to limitations in the completeness and correctness of gold standards, and differences in assumptions and problem formulations. Therefore, we developed five benchmarks to evaluate the quality of the inferred GRNs comprehensively and quantitatively on the aforementioned blood and skin datasets, either based on or independently of gold standards. We focused our comparison on SCENIC, CellOracle, and Dictys given that these methods explicitly model TF binding alongside scRNA-seq data (**Fig. S3a**).

First, in the *TF binding evaluation*, we collected 512 human blood (114 TFs) and 33 mouse skin (16 TFs) ChIP-seq experiments from the Cistrome database^68^ to test whether each method could recover the TF-target gene connections supported by these experiments (**Methods**). In the *TF binding + chromatin loop evaluation*, we further curated the TF binding sites that are more likely to be regulatory by intersecting multiple Erythroid-specific ChIP-seq experiments with chromatin conformation data^53^. We benchmarked each method with Precision-Recall (PR) curves to evaluate method performance at practical GRN sparsity in terms of precision at low recall (**Fig. 4a**). We noticed that full Area Under PR (AUPR) or F1 score primarily covered high recall and failed to capture method performance at low recall (**Fig. S3bc**). This motivated the use of partial AUPR and F_0.1_ scores instead (**Fig. 4bc, Methods**). In all these evaluation metrics, Dictys consistently outperformed existing methods on both datasets (**Fig. 4a-d**).

**Fig 4.**
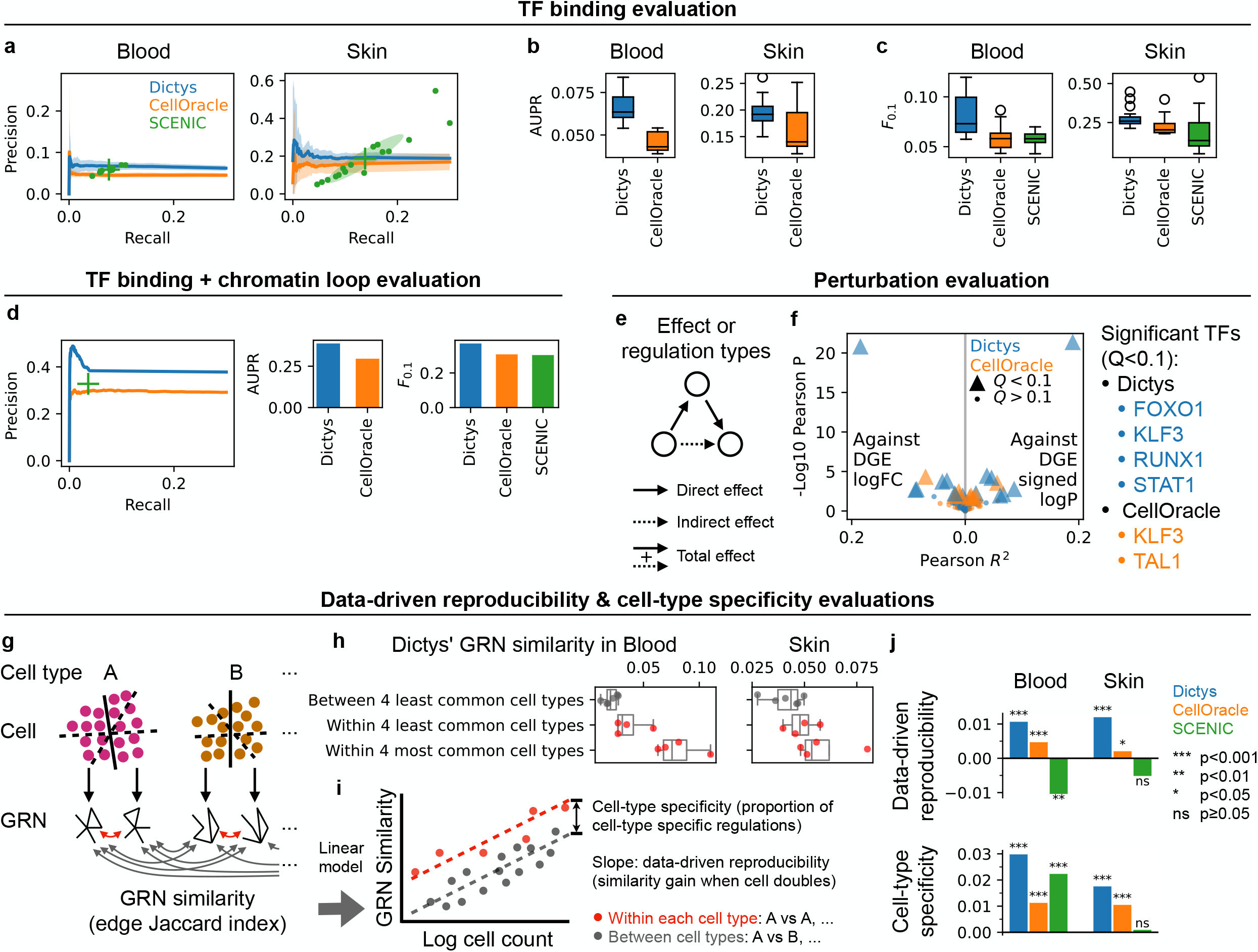
Five quantitative evaluations demonstrated superior performance from Dictys over existing methods. **abc** Precision-Recall curves/dots (**a**), partial AUPRs of **a** (**b**), and F_0.1_ scores (**c**) when comparing cell-type specific GRNs against tissue-specific ChIP-seq experiments. For continuous networks (**a**), PR curve and shade indicate mean and standard deviation across cell types, respectively. For binary networks (**a**), cross, dot, and shade show mean, data point, and covariance across cell types, respectively. For **bc**, each data point is one cell type. **d** Precision-Recall curves/dots (left), partial AUPRs (center), and F_0.1_ scores (right) when comparing Erythroid-specific GRNs against tissue-specific ChIP-seq + Erythroid HiC experiments. **e** A schematic illustration of indirect and total (direct + indirect) effects defined from direct effects on the GRN. **f** Evaluation of total-effect GRN using Pearson test against DGE logFC (left) or signed logP (right) in perturbation studies (single TF KO/KD v.s. control). Each dot represents a cell-type-TF pair while color and shape indicate method and significance, respectively. **g** Schematic plot for GRN similarity quantification for each method within the same cell type (red) and between different cell types (gray) with random cell partitions. **h** Demonstration of different factors affecting GRN similarity – GRN rewiring between cell types and the amount of data (i.e. cell count) – by comparing different cell-type pairs. **i** Schematic plot for the systematic decomposition of GRN similarity contributions from data-driven reproducibility and cell-type specificity with linear model on all cell-type pairs. Dashed lines represent the best linear fit. Colors indicate category of cell-type pair. **j** Dictys consistently had best GRN data-driven reproducibility and cell-type specificity in both datasets.

TF binding does not always lead to regulation of nearby genes, therefore we next developed the *perturbation evaluation* with Knockout (KO) or Knockdown (KD) experiments of hematopoietic TFs from the KnockTF database^69^. To quantify the ability to recapitulate differential gene expression (DGE) statistics (logFC or signed log P-value), we computed the Pearson correlations between these statistics and the predicted downstream effects of in-silico TF perturbations as propagated on the reconstructed continuous networks. To capture indirect effects, we used the total-effect GRN for comparison, either as the steady-state effect of each TF’s transcriptional rate change for Dictys, or with three-step propagation as described for CellOracle (**Fig. 4e, Methods**). Dictys demonstrated superior performance in P-value, R^2^, and the number of significant TFs (Q<0.1, **Fig. 4f, Fig. S3d**). To understand this superior performance, we compared total-effect GRNs against direct and three-step propagated GRNs within Dictys. Total-effect GRN demonstrated clear advantages by accounting for feedback loops and infinite-step propagations with stochastic process network (**Fig. S3d**).

In the *cell-type specificity and data-driven reproducibility evaluations*, we assessed the proportion of cell-type specific and reproducible edges, which together determine the effectiveness of comparative network analyses. By comparing GRNs reconstructed from non-overlapping cell subsets from the same or different cell types (**Fig. 4g, Methods**), we found that GRNs were less similar (in Jaccard index of edge overlap) between cell types than within each cell type. This is expected given that GRNs are rewired in differentiation. In addition, we found that GRNs are more similar with more cells because extra information reduces inference variance (**Fig. 4h**). Consequently, we systematically decomposed GRN similarity with a linear model into contributions from cell count and cell type sameness, as (data-driven) reproducibility and cell-type specificity respectively, emphasizing data-driven discovery capability (**Fig. 4i**). On both datasets, Dictys was more reproducible and cell-type specific than SCENIC and CellOracle (**Fig. 4j, Fig. S3e**).

Finally, we comprehensively investigated how different parameter choices might influence network reconstruction to guide Dictys users (**Fig. S4**). As expected, searching for TF binding sites with peaks rather than footprints could improve recall at the cost of precision. We also tested optional filters of footprint-target gene links using co-accessibility with the transcription start site (TSS)^17^ or using the association with target gene expression^22^, among other parameter settings. Despite a minor overall advantage from the default setting, no alternative setting provided a consistent gain or loss of performance in the TF binding, TF binding + chromatin loop, data-driven reproducibility, and cell-type specificity evaluations.

In conclusion, our systematic set of benchmarks showed that Dictys outperformed existing methods in five different evaluations, while also providing robust and cell-type specific network inference across wide biological applications simulated by various parameter choices.

### Dynamic GRN provides time-resolved discovery and investigation of TFs and regulations in human hematopoiesis

In principle, single-cell technologies enable the study and characterization of continuous processes such as development or disease progression. However, this can be achieved only by computational methods specifically designed to leverage this possibility. Although existing GRN inference methods have been applied to discrete cell groups, Dictys can use any continuous cell ordering (e.g. time, pseudo-time, RNA-velocity, or lineage data) to reconstruct dynamic GRNs and to uncover the continuous rewiring of networks. To showcase this analysis in hematopoiesis^30^, we first inferred pseudo-time trajectories for three developmental lineages (Erythroid, B-cell, and Monocyte) with STREAM^70^ and integrated chromatin accessibility information from matching scATAC-seq with ArchR^71^ (**Fig. 5a, Methods**). Using pseudo-time as a surrogate for developmental order, we inferred one dynamic GRN for each lineage for the (pseudo-)time-resolved discovery and investigation of development-associated TFs and regulations (**Fig. 1e**).

**Fig 5.**
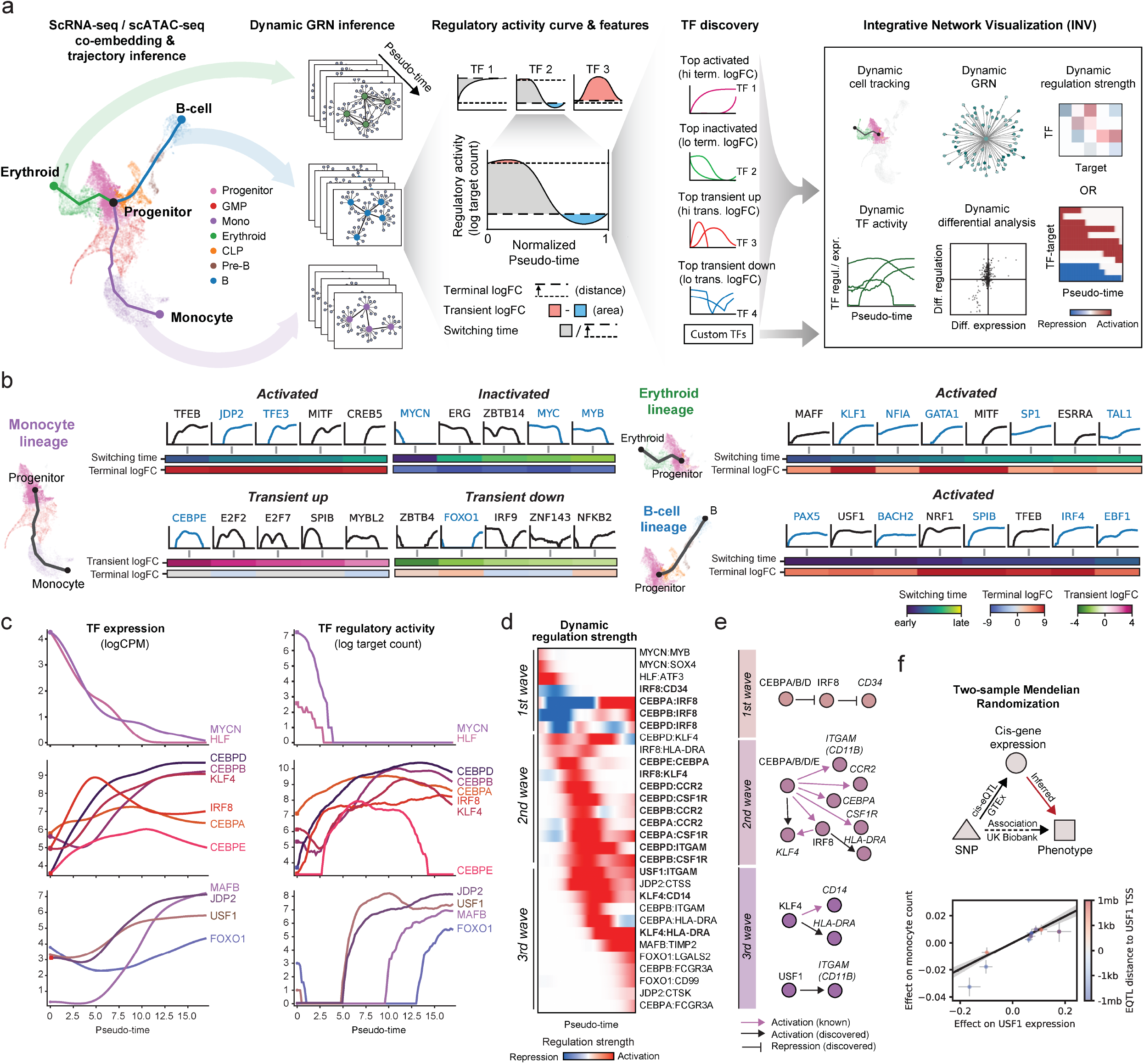
Dynamic GRN reconstruction, TF and regulation discovery, and time-resolved investigation using Dictys. **a** Analytical workflow of dynamic GRN inference, TF discovery from regulatory activity curve, and in-depth investigation with INV for three developmental lineages in human hematopoiesis. **b** Unbiased TF discovery with four distinct patterns (activated, inactivated, and transient up/down) of regulatory activity found established (symbol in blue) and putative (symbol in black) TFs in all three lineages. **c** Expression (left) and regulatory activity (right) curves of prominent TFs during monocyte development. **d** Time-dependent strengths of individual regulations of lineage-defining markers by master TFs in **c**, as grouped in three waves based on when each regulation peaks. Bold regulations are illustrated in **e**. **e** Schematic illustration of prominent regulations recovered in each wave in **d. fg** Two-sample Mendelian Randomization schematics (**f**) and results (**g**) validated and refined TF discovery from dynamic GRN, particularly with the significant effect of USF1 expression on monocyte count (slope). Each dot is a SNP. Shades and error bars indicate standard error.

For each TF, dynamic GRN can directly quantify its time-dependent regulatory activity with a curve. This *regulatory activity curve* contains important information on the potential mechanism of action of a given TF that can be summarized by the curve’s geometric features (e.g. distance and area): terminal logFC, transient logFC, and switching (on/off) time (**Fig. 5a, Fig. S5a, Methods**). Top TFs in terminal or transient logFC unbiasedly captured established driver TFs in activated, inactivated, or transient up/down patterns in each lineage, such as JDP2, TFE3^72,73^ (activated), MYC, MYCN and MYB^74,75^ (inactivated) and CEBPE^36,37^ (transient up) in monocyte; KLF1, GATA1, TAL1, NFIA, SP1^32–35^ (activated) in erythroid; and BACH2, PAX5, SPIB, IRF4, EBF1^39,40,76^ (activated) in B cell (**Fig. 5b**); among other TFs and patterns (**Fig. S5bc, Supplementary Table 4**). These known TFs demonstrate the utility of regulatory activity curves beyond discrete cell types and how our unbiased analysis can prioritize other less known or novel development-associated TFs.

In addition, for these established TFs, Dictys could also delineate their GRN rewiring in sequential waves. As progenitor cells started to differentiate to monocytes, we detected the first wave consisting of Stem/Progenitor-related TFs with declining expression, regulatory activity, and putative individual regulations (HLF-*ATF3*, MYCN-*MYB/SOX4*^31,74,75,77,78^, **Fig. 5cde, Supplementary File 2**). Meanwhile, those of the second wave including myeloid lineage-defining factors, rose steadily and peaked in GMP and early monocytes (CEBPA/B/D-*CCR2/CSF1R/KLF4/ITGAM*^36,37,79–81^). The third wave comprised monocyte lineage-defining TFs and their regulations (MAFB-*TIMP2*, FOXO1-*CD99/LGALS2*, KLF4-*HLA-DRA/CD14*, JDP2-*CTSS/CTSK*^38,73,81–87^). Dictys also inferred potential multi-modal regulations throughout these waves with time-dependent directionality shifts, such as CEBPA/B/D-*IRF8* that was initially repressive but later turned activating. These time-dependent multi-modal effects may provide novel insights on the regulatory cascade in development, e.g. of the stem cell marker CD34^88^ and the major monocyte driver/marker *KLF4* and *HLA-DRA*^36,81^.

To refine TF discoveries with orthogonal information, we performed two-sample Mendelian Randomization^89^ (MR) to identify TFs whose mRNA expression may affect monocyte development (count or proportion). We used cis-expression quantitative loci (cis-eQTLs) of each TF from the GTEx project^90^ as instrumental variables and considered their causal effects on these organismal phenotypes from UK Biobank^91^ (**Fig. 5f, Methods**). Among the TFs suggested causal to monocyte development by MR, USF1 was within the top 10 activated TFs, more differentially regulated than differentially expressed, and also previously linked to atherosclerosis and inflammation^92,93^. Our data-driven approach can discover potential novel TFs whose accuracy can be further improved with orthogonal analyses.

On the Erythroid and B cell lineages, their dynamic GRNs were likewise productive for time-resolved discoveries and investigations of established and novel TFs and regulations (**Figs. S5, S6, Supplementary Files 1, 3, 4**). Notably, FOXO1 is known to carry distinct functions at different stages of B cell development^94^, as reflected by its irregular regulatory activity curve. In contrary, this signature was unobservable in its monotonic expression level curve. Overall, Dictys’ dynamic GRN reconstruction provided fine time resolution and regulatory insights into hematopoiesis not available from coarse-grained cell clusters or mean expression-based analyses.

## Discussion

In this manuscript, we describe Dictys, a GRN inference method to reconstruct, analyze, and visualize context specific and dynamic GRNs from single-cell transcriptome and chromatin accessibility data. Dictys leverages context specific TF footprinting to prioritize direct regulation through DNA binding, stochastic process models to allow for feedback loops, probabilistic programming to account for scRNA-seq technical variation, and dynamic network reconstruction for time-resolved TF activity and regulation. Our comprehensive benchmarks showed that Dictys has superior performances against existing static GRN inference methods, especially in cell-type specificity and data-driven reproducibility. Dictys enables comparative GRN analyses from discovery to investigation, from gene to regulation, and from discrete groups to continuous processes. Importantly, Dictys uncovered biological insights in blood and skin systems not captured by current single-cell analyses based on mean expression changes and/or cell clusters. Dictys is a versatile tool for a wide range of GRN studies and application scenarios from inference to analysis and visualization.

GRN inference is a long-standing fundamental question in biology with many challenges. Dictys extends steady-state linear models of GRN to stochastic process models with steady-state observations. However, our method does not consider nonlinearity, non-steady-state, or mechanistic models. Neither does it incorporate other data modalities that may improve inference, e.g. unspliced mRNAs, proteins, DNA methylation, single-cell perturbations, or cell lineage measurements. Dictys assumes steady states but still uncovers rich biology in non-steady-state developmental systems. Unsurprisingly, GRN inference has high variance when cell count is limited, which we stabilize with kernel smoothing and target count based analyses. TF binding prediction accuracy can be limited by cell count and by the lack of direct single-cell measurements from biochemical/chromatin binding assays. Evaluation of GRN inference methods remains difficult with limited quality and quantity of gold standards. We believe these are exciting directions for future work.

Single-cell technologies have made comparative GRN inference and analysis feasible for individual samples and across multiple modalities. Dynamic GRN inference methods like Dictys can uncover cell circuits and quantify gene regulations at high throughput and resolution, providing a much broader scope than individual gene expression for future biological discovery and investigation.

## Acknowledgements

We wish to thank Jason Buenrostro, Huidong Chen, Yan Hu, Vinay Kartha, and Qian Qin for helpful discussions. L.P. received support from the U.S. NIH (R35 HG010717). We thank the Pinello lab members for helpful discussions.

## Methods

### Dictys

#### Transcriptome quality control (QC)

On the transcriptome data of selected cell subpopulation, we removed cells with <200 total reads or <100 expressed (with >0 reads) genes, and genes with <50 total reads or expressed in <10 cells. This was done iteratively until no gene or cell was removed. For joint measurement of transcriptome and chromatin accessibility, cells were removed from both modality profiles.

#### TF binding network

For each cell subpopulation, bam files were first aggregated to pseudo-bulk after removal of chrM and chrUn, and processed with macs2 (2.2.7.1) to obtain chromatin accessibility peaks (top 500k by default). ENCODE blacklisted genome regions were removed^95^. We then used wellington_bootstrap.py^96^ (pyDNase 0.3.0) to annotate (top 100k by default) TF footprints, and Homer (4.11) findMotifsGenome.pl function to scan the TF footprint regions for known TF motifs from existing databases (see below).

To score the TF links to their potential target genes, we created a multipartite graph with the above outputs from each TF to its known motifs, from each motif to its containing footprints, and from each footprint to its nearby genes (up to 500kb from TSS by default). Each motif-footprint link was scored additively based on the wellington footprint probability score *W*, Homer motif purity score *H*, and the distance between footprint and target gene TSS *D* as log_10_ *W* + log_10_ *H* − 10^−6^*D* ^13^. The maximum value over all directed paths was used for each TF-target link on the multipartite graph. This formed a continuous TF-binding network with TF footprinting from chromatin accessibility profile. Each gene’s top (20 by default) putative regulator TFs were then selected to form the binary TF-binding network to constrain downstream network reconstruction to reduce search space.

#### Stochastic kinetic model for transcriptome

We characterized the transcriptomic kinetics of each cell with the stochastic process following an empirical linear model as a stochastic differential equation (SDE)

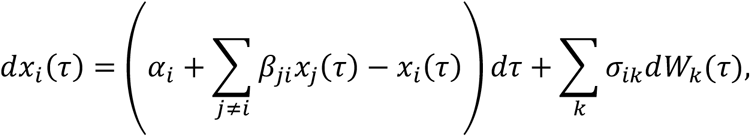

where *x*_*i*_(*τ*) is log expression level of gene *i* at hypothetical time *τ* to account for the multiplicative nature of stochasticity and chemical master equation, *α*_*i*_ is the empirical intercept representing the steady-state net *basal transcription rate* for gene *i, β*_*ji*_ is the transcriptional regulation strength from gene *j* to gene *i* and also the *stochastic process GRN* to be reconstructed, −*x*_*i*_(*τ*) fixes the time scale of this equation with the cell’s overall transcriptional capacity as a *regularization* term, *dW*_*k*_(*τ*) is the Wiener process to represent independent modes of intra- and extra-cellular *stochasticity*, and *σ*_*ik*_ is their influence on each gene. Only the *β*_*ji*_ entries allowed by the TF-binding network can have non-zero values. This SDE describes the multi-variate Ornstein-Uhlenbeck process whose analytical steady-state solution for each cell at *τ* → *∞* follows the multi-variate normal (MVN) distribution in matrix form for each cell *k* as

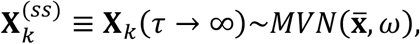

where 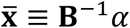 is the mean expression vector for genes, **B** ≡ **I** − *β*^*T*^, and the covariance matrix *ω* is the solution of the continuous Lyapunov equation

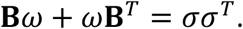

This fully determines the steady-state distribution of gene expression given the stochastic process parameters.

See **Supplementary File 1** for details.

#### Relative gene abundance

For each cell *k*, we modelled the true gene expression vector 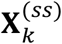 as an independent sample from the above MVN distribution. The relative gene abundance 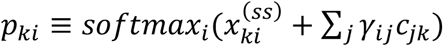 for gene *i* contains contributions from the true expression 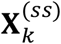 and known confounders. Here *c*_*jk*_ is the quantification of confounder *j* on cell *k*, and *γ*_*ij*_ is the effect size of confounder *j* on gene *i*. Because whether each confounder’s effects propagate through the GRN is regarded unknown, encoding confounder effects outside the GRN can account for both types of confounders universally. We did not include any technical confounders although they were supported by the model, because batch effects were not strong in low dimensions according to the original publications. Finally, the softmax function converts expression from log to linear scale and from arbitrary to relative scale.

#### Probabilistic modelling and parameter estimation

The final UMI read count of each gene *i* as *y*_*ki*_ then follows the binomial distribution *B*(*n*_*k*_, *p*_*ki*_), where *n*_*k*_ ≡ Σ_*i*_ *y*_*ki*_ is the total read count for cell *k*.

In total, this is a probabilistic generative model for the UMI count matrix with stochastic process parameters *α, β, σ*, and confounder effect *γ* (**Supplementary File 1**). We reparameterized *α* with 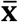 to directly reflect steady state mean expression. Only entries in *β* allowed by the TF-binding network can have non-zero values. We used a low-rank parameterization for *σ*, equivalent with a low-rank MVN distribution for gene expression stochasticity, as *σ* ≡ (*σ*^(*v*)^, *σ*^(*cov*)^). Here *σ*^(*v*)^ is a diagonal matrix to model independent stochasticity for each gene, and *σ*^(*cov*)^ models hidden confounders that apply stochasticity with covariance in low rank form (by column). We tested different numbers of hidden confounders and used 0 by default.

Altogether, this model contains parameters 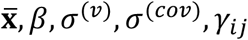. Parameter estimation with maximum likelihood estimators was achieved through training the probabilistic model with Pyro (1.6.0) with initial learning rate 0.01 and learning rate decay 0.999 for 4000 iterations against the observed UMI count matrix **Y**.

#### Steady-state total-effect network

The raw steady-state total (direct + indirect) effect network 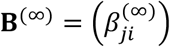 was defined as the consequence of perturbation of the basal transcription rate *α* on the expected steady-state expression of other genes 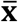. Specifically, the raw steady-state total-effect network 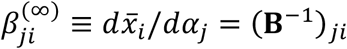, and therefore **B**^(*∞*)^ = **B**^−1^. For comparison with DE in KO/KD experiments, the raw total-effect network was further scale-normalized for each TF by its effect on the total absolute mRNA abundance to account for library size normalization.

#### GPU acceleration

To achieve GPU acceleration without a GPU-based Lyapunov equation solver for *ω*, we replaced the computation of *ω* with a mean-squared loss term between the left- and right-hand sides of the Lyapunov equation (**Supplementary File 1**). We assigned a large constant multiplier as a hyperparameter to this extra loss to ensure deviations are minimal. This design allows simultaneous estimations of model parameters and the approximate solution for Lyapunov equation.

#### Post processing

The network edge strength *β* is linearly proportional to the square root expression variance of the target gene and inversely proportional to that of the regulator. However, variance estimation is challenging and often negatively biased in scRNA-seq datasets due to sparsity^20^. To avoid bias propagation to network edge strength, we defined a relative edge strength by scale-normalizing *β* with stochasticity-induced variance, as

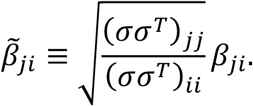

Therefore, 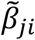 captures the relative amount of variance in target gene expression affected by TF expression and no longer depends on the overall variance scale of TF or target. It is the final output of edge strength in individual GRN inference from a given cell subpopulation.

#### Moving window in dynamic GRN inference

Moving window was constructed from the projected cell locations onto the trajectory by shortest distance on low dimensions. The trajectory should be provided by user and can be inferred with any software and from any source of information. Here we used STREAM for trajectory inference. See “Single-cell datasets”.

Moving window was defined as a set of (overlapping) fixed-sized cell subpopulations on the trajectory. The construction of moving window was subject to two constraints: a hard lower bound on the number of overlapping cells between neighboring subpopulations, and a soft upper bound on the pseudo-time distance between neighboring subpopulations. The pseudo-time distance was defined as the cross-subpopulation mean cell distance subtracted by the maximum (over two populations) of in-subpopulation mean cell distance. The mean cell distance was the average trajectory distance between all cell pairs. Trajectory distance was the low-dimensional distance between points on the trajectory computed along trajectory edges.

To construct a moving window for RNA profiled cells, one subpopulation was created for each trajectory node and for each cell as an anchor, both with the same number of nearest cells based on trajectory distance. Then, each anchor-cell associated subpopulation was checked once at a random order and removed only if neither of the above constraints would be violated by its two neighboring subpopulations. All remaining subpopulations formed the moving window along with their neighborhood relations. For non-joint ATAC profiled cells, (the same number of) nearest cells were selected for each remaining anchor cell.

#### Gaussian kernel smoothing

To obtain the value of a “dynamic variable” at any pseudo-time point on a particular differentiation trajectory from finite subpopulations, we interpolated the variable’s known values from nearby subpopulations with an exponentially decaying weight 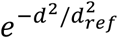, where *d* is the pseudo-time distance to each subpopulation’s center and *d*_*ref*_ = 1.5 pseudo-time units. Subpopulations with anchor cells on a different branch of the trajectory of interest were ignored for kernel smoothing.

Dynamic variables that received direct Gaussian kernel smoothing from their known values in the subpopulation include gene expression (pseudo-bulk) CPM and each edge’s strength in the GRN. Dynamic GRN is the collection of dynamic edge strengths. All their downstream variables were consequentially affected, such as number of target genes, differential analysis logFC, regulatory activity curve.

### Network analysis in Dictys

#### Network binarization

Many established network analysis approaches do not accept edge weights. Therefore, continuous networks from Dictys were first converted to binary networks with the provided sparsity rate, i.e. the proportion of edges that are positive. Lower sparsity rate selects more confident and stronger gene regulations based on the absolute value of edge strength. On the experimental perspective, this binarization approach can identify the strongest edges for further validation and relies solely on the ranking of edge strengths.

Many factors vary from biological to technical aspects, such as cell type, TF activity, motif length, and data quality. Therefore, the ideal sparsity to illustrate and understand biology can also differ greatly. There is no consensus on how to choose network sparsity, so we leave it to the user.

Throughout this study, we used the constant sparsity rate = 0.01 (1% of all edges positive) as a rule of thumb and to avoid data over-fitting and over-interpretation. The only exception is subnetwork visualization for TFs with too few strongly detected regulators or targets (i.e. GATA1, HLTF, Dlx3), where we increased the sparsity rate to 0.05 to include more genes.

#### Expression and regulation specificity and regulation marker genes

For each cell type *j*, TF *i*’s (absolute) *outdegree centrality* (i.e. regulatory activity or target count) were first computed from the binary network as *centrality*_*ij*_. The *relative outdegree* normalized the TF’s regulatory activity in context, as 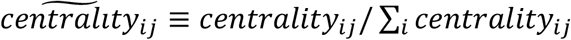. TF *i*’s regulation specificity to one cell type characterized the probability of its randomly picked regulation coming from this cell type, as 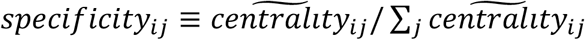. TF *i*’s overall regulation specificity was the relative difference between the entropies from its specificity to each cell type and from uniform distribution, as *specificity*_*i*_ ≡ 1 + ∑_*j*_ *specificity*_*ij*_ log *specificity*_*ij*_ / log #_*j*_.

For direct comparison, expression specificity was a repetition of above after replacing outdegree with CPM and represented random draws of RNA reads instead of regulatory relations. In this study, we selected regulation marker genes as those with overall specificity ≥ 0.5 and with specificity ≥ 0.4 and outdegree ≥ 20 in at least one cell type. For each cell type, we visualized the top (up to) 10 regulation markers based on specificity.

Therefore, expression and regulation marker gene discovery depends on the granularity of cell type annotations. When one cell type is divided into subtypes more finely than others (e.g. different T cell subtypes in the blood dataset here), each subtype may recover fewer marker genes due to lower cell count and reduced specificity when compared with other subtypes.

#### Differential regulation and differential expression logFCs

For direct comparability, we defined the logFC as the ratio of log_2_(*x* + 1) between two cell types, where, x is regulatory activity (i.e. target count) for differential regulation and CPM for differential expression. We also used logCPM and log target count to indicate log_2_(*CPM* + 1) and log_2_(*target court* + 1) in this paper.

#### Context specific TF ranking

TFs were directly ranked for context (e.g. cell-type) specificity from the logFCs in differential expression or differential regulation. The combined ranking was obtained from the mean logFC to integrate expression and regulation specificity.

#### Geometric characteristics/features of TF regulatory activity curve from dynamic GRN

Regulatory activity 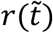 as a function of normalized pseudo-time 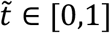 was defined as log (target count +1) for the dynamic GRN, after binarization at each pseudo-time point. We defined three intuitive characteristics of TF regulatory activity 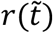 from geometric measures including distance (or difference) and area under the curve. Terminal logFC = *r*(1) − *r*(0) is the regulatory activity difference (also differential regulation logFC) between the final and initial pseudo-time points. Transient 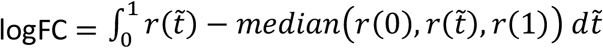 is the net area of regulatory activity above or below its both initial and final values. 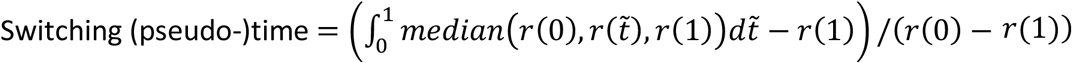 is the normalized area between current and final regulatory activity within initial and final bounds. These integrals were computed with evenly distributed pseudo-time points (by default 100 time points) from beginning to end of the lineage.

#### GRN subnetwork visualization

Subnetwork visualization was performed with force-directed layout with Cytoscape (3.5) for context specific GRN or Dictys (adapted from networkx) for dynamic GRN, with the weight of each edge proportional to the absolute value of regulation strength.

### Data

#### Single-cell datasets

Single-cell data were retrieved from previous studies, including the human blood dataset from separate scRNA-seq and scATAC-seq experiments of total and CD34(+)-enriched bone marrow mononuclear cells from healthy donors^30^ and the mouse skin dataset from the SHARE-seq experiment^22^.

Cell annotations such as cell type and low dimensional embedding coordinates were obtained from the original studies. For the human blood dataset, mapping cells from scATAC-seq to cells from scRNA-seq was achieved with ArchR^71^ (0.9.5, predicted cell) following its tutorial (https://www.archrproject.com/bookdown/index.html). Trajectory inference was achieved with STREAM^70^ (1.0) with epg_alpha=0.05, epg_mu=0.01, epg_lambda=0.2, and epg_ext_par=1.

#### ChIP-seq datasets

For original ChIP-seq results (for TF binding evaluation), we obtained 512 human blood (114 TFs) and 33 mouse skin (16 TFs) ChIP-seq experiments from the Cistrome database^68^. For ChIP-seq + HiC results in Erythroid (for cell-type specific GRN analysis and TF binding + chromatin loop evaluation), we further combined different experiments in Erythroid for each TF using bedtools (2.9.2) to obtain a consensus of binding sites (peaks found in at least 2 experiments), which was only available for 16 TFs (ATRX, BPTF, GATA1, GATA2, HDAC1, HDAC2, KLF1, LDB1, NCOA4, NFE2, NR2C2, POLR2A, SETD1A, STAG1, TAL1, ZBTB7A).

Peaks were mapped to the nearest genes when gene-level analysis was needed. For ChIP-seq + HiC results, peak-gene mappings were removed if there was no chromatin interaction loop in promoter capture Hi-C in Erythroid^53^ between the peak and any peak in the promoter of the target gene (+/-10bp of TSS).

#### TF motif database

We used the Homer-formatted (p-value 0.0001), full human or mouse collections from the HOCOMOCO (v11) database^97^.

#### Optional filters on footprint-target gene links

To filter footprint-target gene links with co-accessibility with TSS, we ran Cicero^98^ (1.3.4.10) on the peaks of cell-type specific pseudo-bulk chromatin accessibility profiles, and otherwise followed CellOracle’s approach^17^. Footprint-target gene links were removed if the peak containing the footprint had co-accessibility score <0.54 with all the peaks around the TSS of target gene (distance <5kb). See “Existing methods”.

To filter footprint-target gene links with association with target gene expression in mouse skin, we downloaded these associations computed at cell population level from the Supplementary Table 4 of ^22^. Footprint-target gene links were removed if the peak containing the footprint had association p-value >0.1 with target gene expression.

### Cell-type specific GRN analyses

#### TF footprint quality assessment

TF footprint meta-plots (average Tn5 signal) were produced using wellington (dnase_average_profile.py -A, pyDNase 0.3.0). TF footprint saturation plot was produced by counting the number of footprints in random cells subsampled from erythroid subpopulation for 10 times.

#### Gene enrichment analysis

Gene enrichment analysis was done with EnrichR^72^ (March 2021 update) using all genes as a background gene set (default).

#### Integrative Genome Viewer tracks

ATAC-seq signal tracks were reconstructed from bam files using Deeptools (4.1.3) with command bamCoverage --binSize 50 -e 200 --normalizeUsing RPKM. Integrative Genome Viewer (2.10.0) was used to visualize epigenomic data on genomic loci of interest.

### Evaluations

#### Existing methods

For SCENIC, we used pyscenic (0.10.3) following the official tutorial (https://pyscenic.readthedocs.io/en/latest/tutorial.html). Feather files for “500bpUP” and “TSS+/- 10kbp” were used.

For CellOracle (0.6.4), we followed the official tutorial (https://morris-lab.github.io/CellOracle.documentation/tutorials/index.html). The tutorial used a co-accessibility (with TSS) cutoff 0.8 to filter the associated peaks for each gene, which however yielded few network edges. We performed a cutoff scan with 0.01 step size. We determined that 0.54 could give the most similar filtering rate of associated peaks and should be used. All genes, instead of top 2000 highly variable genes, were used to enable benchmarking with other methods.

#### TF binding evaluation

We obtained tissue-specific (blood or skin) ChIP-seq peak data from the Cistrome database. For each experiment (by GSMID), the top 100 peaks (by peak score) were regarded as positive hits and annotated to the nearest gene. The TF binding gold standard was defined as TF-target pairs that were positive hits in at least one of the experiments.

Precision-recall curves (for continuous GRNs by Dictys and CellOracle) and scores (for binary GRNs by SCENIC) were computed for each cell-type specific GRN against the above tissue-specific gold standard. Only the overlapping TFs/target genes showing up in the cell-type specific GRN and having at least one target/regulator in the gold standard were considered.

*F*_0.1_ score is defined as

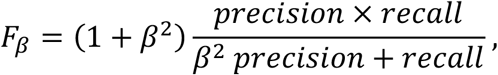

for *β* = 0.1, which is similar to *F*_1_ score for *β* = 1 but focuses on precision at low recall, effectively giving precision 10x importance than recall^99^. Performance at large recall was not investigated because such weak cutoffs would 1) obtain overly dense GRNs especially for those driven by TF binding (e.g. each TF has 4000 direct targets on average), and therefore are rarely used in practice and 2) be incomparable with SCENIC’s sparsity. The *F*_0.1_ scores for continuous GRNs were computed as its largest value across all possible cutoffs that converts a continuous GRN to a binary GRN.

For the same reason, only partial (recall<0.3 in agreement with PR curve figures) AUPRs were used to summarize the PR curves of continuous GRNs (CellOracle, Dictys, and Dictys with alternative parameters) for evaluation.

#### TF binding + chromatin loop evaluation

The TF binding + chromatin loop evaluation was performed against the ChIP-seq + HiC results in Erythroid (intersection of multiple Erythroid-specific ChIP-seq experiments and chromatin conformation data), among overlapping TFs but for all potential target genes, and was otherwise identical with the TF binding evaluation.

#### Total-effect GRN

The total-effect GRN for Dictys was the steady-state GRN. The total-effect GRN for CellOracle was obtained following its “signal propagation within the GRN” approach with three iterations or propagation steps^17^. For benchmarking purposes only, we also utilized the three-step propagation from CellOracle on Dictys’ direct GRN.

#### Perturbation evaluation

We obtained tissue-specific (“haematopoietic and lymphoid tissue”) DGE logFCs and P-values between TF KO/KD and control conditions from the KnockTF database. For each dataset in KnockTF, Pearson correlation test was performed between DGE logFC or signed log P-value vector and the edge strength vector from the perturbed TF to candidate target genes in each cell-type specific total-effect GRN. Only sufficiently strong candidate target genes in the total-effect GRN (inferred edge strength > 0.01, i.e. one unit change in the perturbed TF expression would lead to >0.01 unit change in candidate target gene expression) were included for Pearson test. Pearson tests with <10, 20, or 50 candidate target genes were skipped for robustness. We used these three thresholds of candidate target gene count in parallel, independent evaluations to demonstrate result stability. Q values were computed from P values of the performed Pearson tests with Benjamini-Hochberg procedure. Only methods that infer continuous networks (Dictys and CellOracle) were evaluated because binary networks do not contain directionality or effect size to be propagated on the network, or to be evaluated with Pearson correlation correctly. Within Dictys, evaluation of the direct effect GRN, total-effect steady-state GRN, and CellOracle’s three-step propagation of direct effect GRN also followed the same procedure.

#### Data-driven reproducibility and cell-type specificity evaluations

For Dictys and SCENIC, cells of each cell type were independently randomly split *in silico* into two equal subsets for 10 times to obtain subsampled data (read counts for transcriptome and individual reads for chromatin accessibility) before any preprocessing. A binary GRN was reconstructed for each cell subset (see network binarization), totaling 20 GRNs for each cell type. For CellOracle, since it was designed for whole-population chromatin accessibility profiles, all cells were randomly split *in silico* into two equal subsets for 10 times instead. Then a binary GRN was reconstructed for each cell type in each cell subset, totaling 20 GRNs for each cell type.

GRN similarity was quantified with Jaccard index of the inferred binary edges from non-overlapping cell subsets, i.e. between two subsets in the same split for GRNs of the same cell type (totaling 10 values), or between any two subsets for GRNs of different cell types (totaling 400 values). This could avoid overestimated similarity due to overlapping input data. Jaccard index computation was restricted to edges whose regulator and target genes were both present (i.e. not excluded) in both GRN compared. This could separate node (gene) detection from edge (regulation) detection in GRN similarity.

For each GRN inference method, the mean GRN similarity was computed for each cell type pair, only if both cell subsets had at least 300 cells for blood dataset or 600 for skin dataset to account for different sequencing depths. We then used a linear model to characterize the different sources of GRN similarity between cell types *i* and *j*:

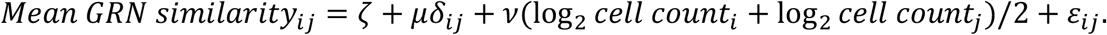

Here *ζ* is the intercept, *µ* is the contribution from cell-type specificity (i.e. proportion of cell-type specific regulations), *δ*_*ij*_ is the Kronecker delta function, *v* is the contribution from data (i.e. data-driven reproducibility or reproducibility gain when cell count doubles), and *ε*_*ij*_ ∼ *i. i. d N*(0, *σ*^2^) is the error. Using *v* instead of *ζ* allows to quantify reproducibility from data instead of model assumptions, e.g. prior knowledge, and therefore can capture the reproducibility of data-driven discovery of novel biology.

Parameter estimation with maximum likelihood, and hypothesis testing with likelihood ratio (for separate null hypotheses: *µ, v* = 0) for the above linear model were performed with Normalisr (1.0.0) based on mean GRN similarity across all cell type pairs.

### Dynamic GRN analyses

#### Two-sample Mendelian Randomization

We performed two-sample Mendelian Randomization with package TwoSampleMR^89^ (0.5.6, Inverse variance weighted) following the guide (https://mrcieu.github.io/TwoSampleMR/articles/index.html) on GTEx (whole blood) cis-eQTL data^90^ and UK Biobank GWAS results^91^. Results were shown with LD clumping for *R*^2^ > 0.2, but other options (no clumping, clumping for *R*^2^ > 0.05 or *R*^2^ > 0.01) gave consistent effect size estimations. UK Biobank GWAS results were restricted to inverse-rank normal transformed traits on both sexes, restricted to monocyte count and monocyte proportion. Two-sample MR was not performed for B or Erythroid cell specific traits because whole blood gene expression was not representative of the actual gene expression levels in these cell types.

## Code availability

Dictys is publicly available at https://github.com/pinellolab/dictys. All the analyses presented in this manuscript were produced with Dictys version 0.1.0 deposited in Zenodo^100^.

## Data availability

The human blood dataset included bone marrow mononuclear cells from healthy donors and was downloaded from GSE139369. The mouse skin dataset was downloaded from GSE140203. The reconstructed cell-type specific and dynamic GRNs and the tutorial data for Dictys are available at ^101^.

## Author contributions

LW, NT, and LP conceived the study. NT developed TF binding network functions. LW developed stochastic process model, network analysis, and dynamic network. GD developed CellOracle benchmarking interface. MH developed dynamic network layout. NT, LW, and TW analyzed and interpreted biological data. LW and NT performed benchmarking. LP and DB supervised the study and provided funding. LW, NT, and LP wrote the manuscript with inputs from all authors.

---

**Fig S1.**
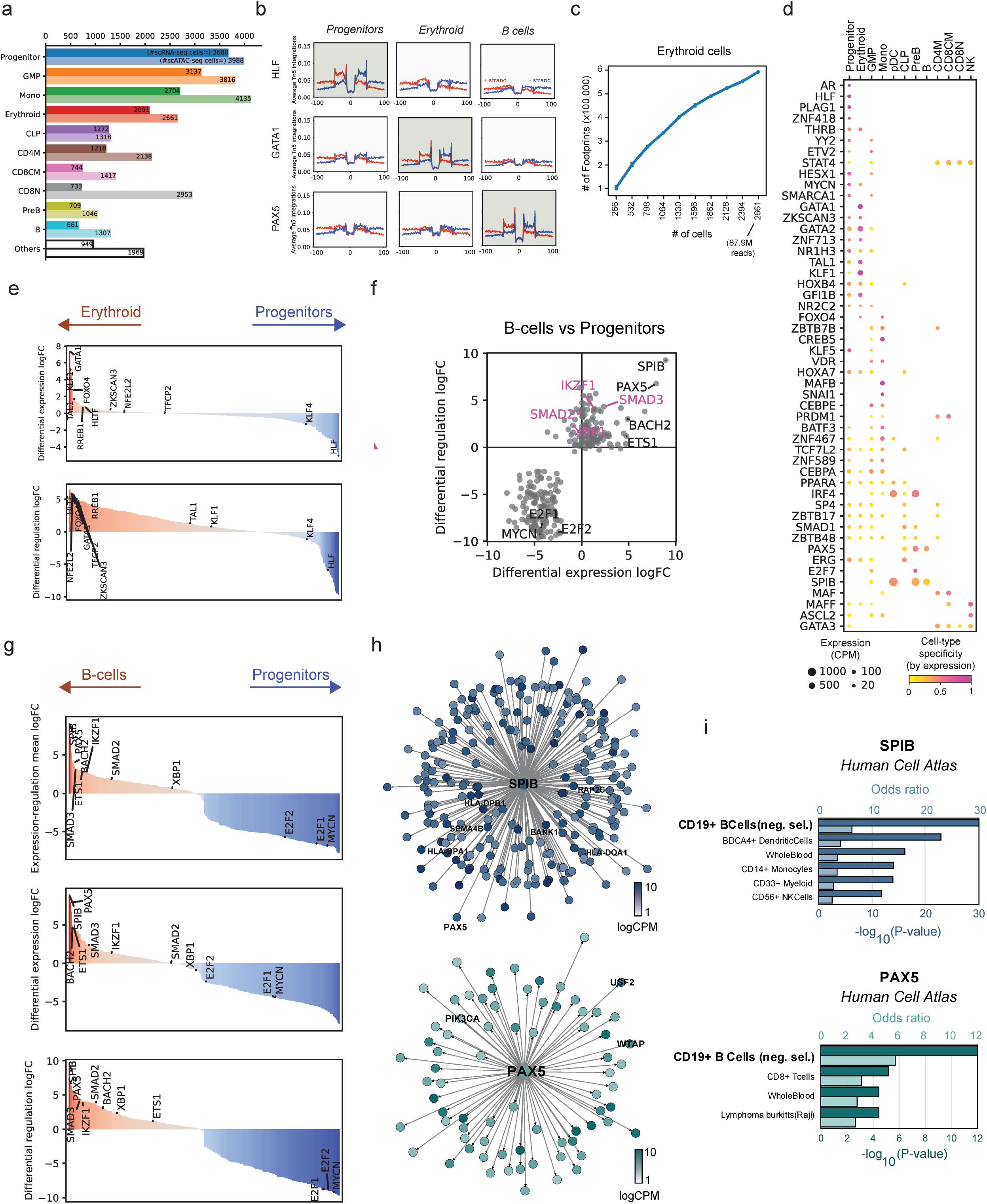
Reconstructing cell-type specific GRNs in human hematopoiesis using Dictys. **a** Bar chart for cell type composition after QC. **b** DNA footprinting recovered binding events of major TFs in their relevant cell types. Total read count of chromatin accessibility (Y) shows a drop at each base (X) towards the discovered binding sites. **c** Footprint discovery saturation plot. Number of footprints detected as a function of the number of cells subsampled from the Erythroid cell population. Mean and standard deviations are shown. **d** Dot plot illustration of expression level and its cell-type specificity for regulation marker TFs discovered in **Fig. 2b**. **e** TF ranking based on logFC of differential expression (top) or differential regulation (bottom) alone between Erythroid and Progenitor cells. **f** Scatter plot for differential expression (X) and differential regulation (Y) logFCs between B cells and Progenitors. **g** Integrative TF ranking (top) and separate TF rankings based on the logFC of differential expression (middle) or differential regulation (bottom) between B cells and Progenitors. **h** Subnetwork plots of inferred targets for SPIB (top) and PAX5 (bottom). **i** Enrichment analysis of SPIB (top) and PAX5 (bottom) targets in **h**.

**Fig S2.**
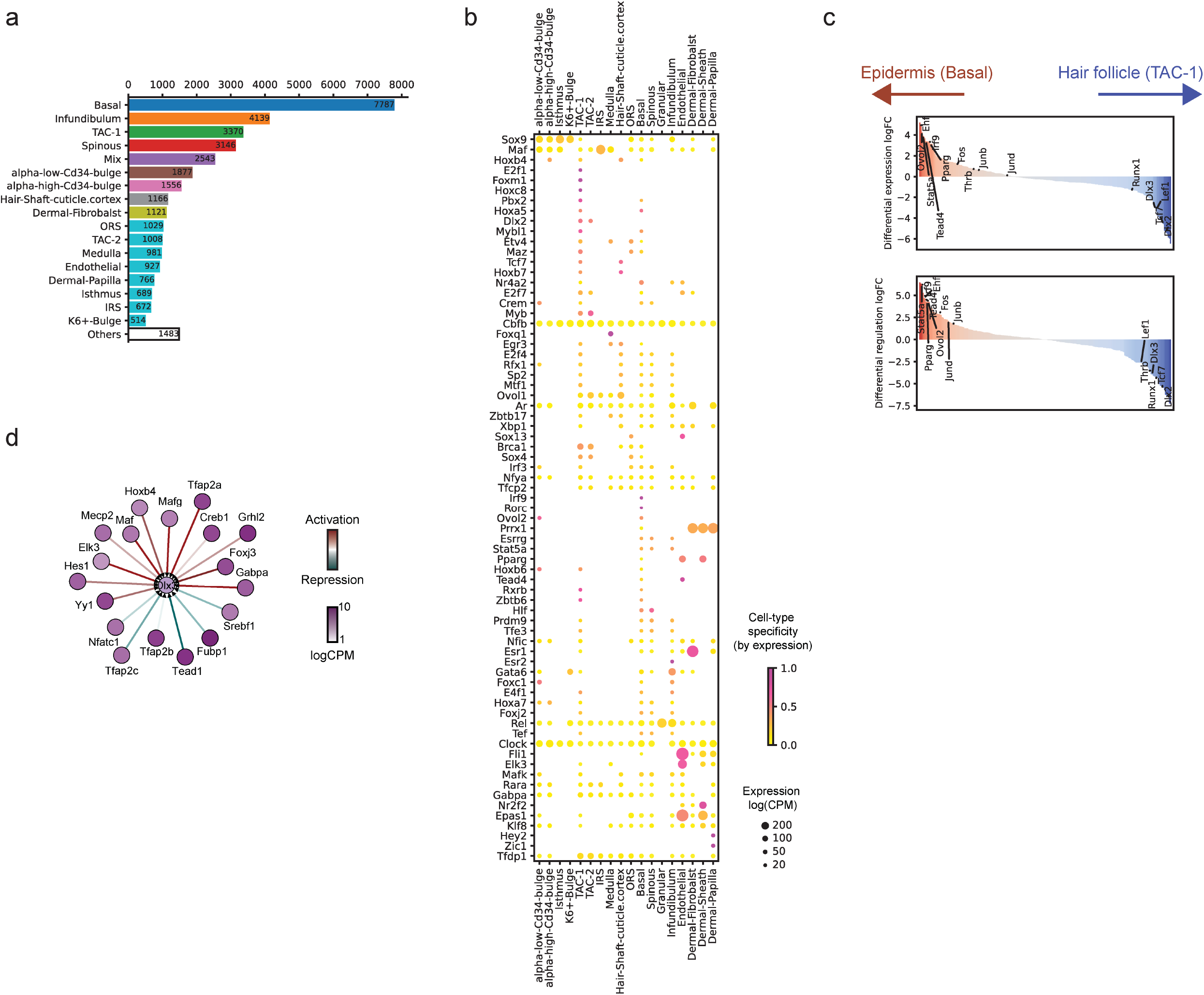
Reconstructing cell-type specific GRNs in mouse skin using Dictys. **a** Bar chart for cell type composition after QC. **b** Dot plot illustration of expression level and its cell-type specificity for regulation marker TFs discovered in **Fig. 3b**. **c** TF ranking based on logFC of differential expression (top) or differential regulation (bottom) alone between epidermis and hair follicle cells. **d** Subnetwork plot of the inferred regulator TFs of *Dlx3* in TAC-1 when the optional filter from transcriptome-chromatin accessibility association was not applied.

**Fig S3.**
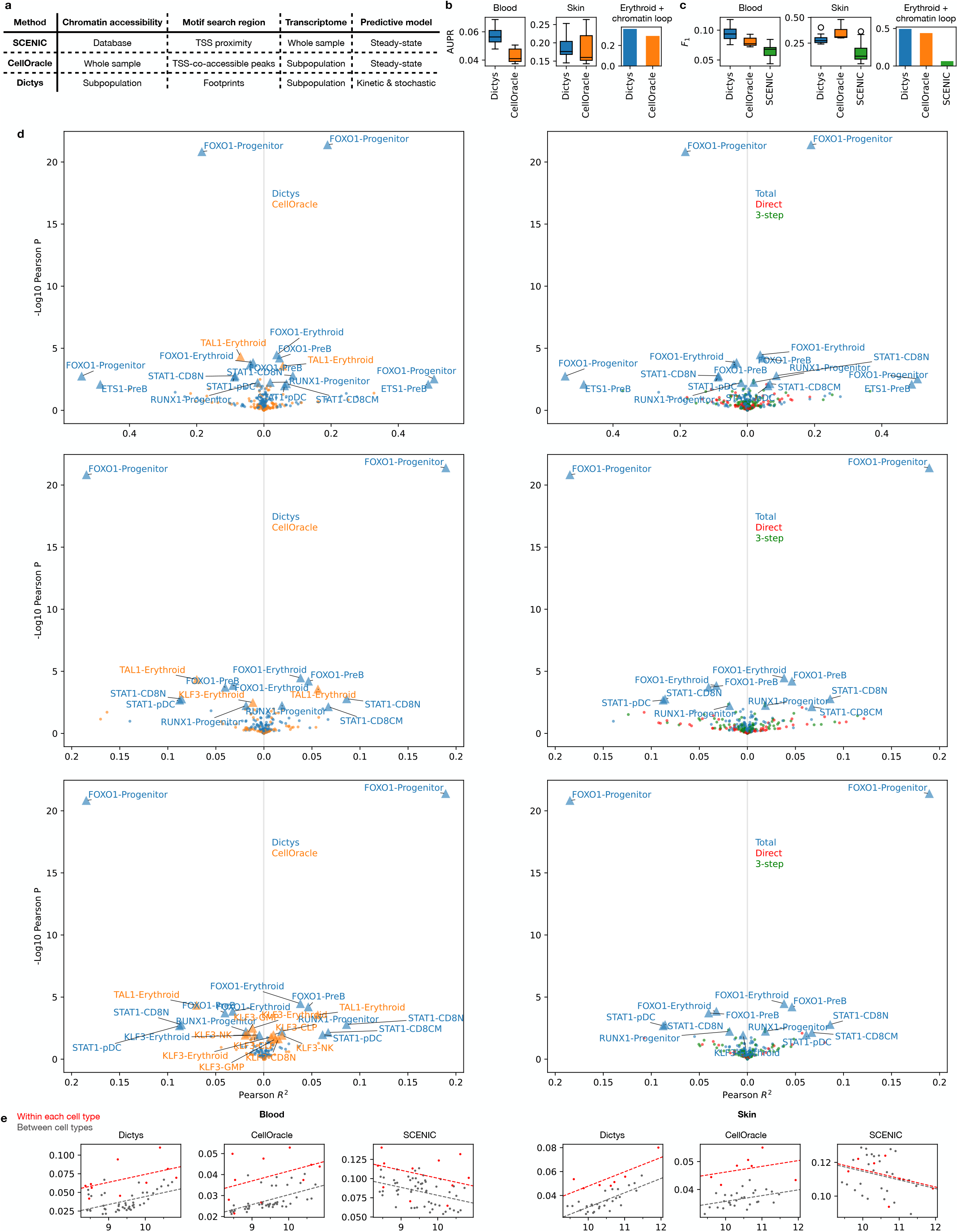
Benchmarking of static GRN reconstruction methods. **a** A table of benchmarked single-cell GRN inference methods that also use TF binding information, with their technical differences. **bc** Full AUPR (**b**) or F_1_ score (**c**) could not capture method performance at low recall in PR curves so well as partial AUPR or F_0.1_ score, for TF binding evaluation (blood, skin) and TF binding + chromatin loop evaluation (Erythroid + chromatin loop). **d** Annotated version scatter plots for perturbation evaluation between CellOracle’ 3-step propagated GRN and Dictys’ total-effect GRN (left) or between Dictys’ total-effect, 3-step as in CellOracle, and direct networks (right). Total-effect GRN from Dictys could recapitulate TF KO/KD results best. Evaluation results were robust at different thresholds (10, 20, or 50, top to bottom) for the minimum number of target genes for each TF to be included in the evaluation. **Fig. 4f** is an unannotated version of the bottom left panel. **e** Demonstration of linear fit (**Fig. 4i**) for Dictys (default settings), CellOracle, and SCENIC on human blood and mouse skin datasets.

**Fig S4.**
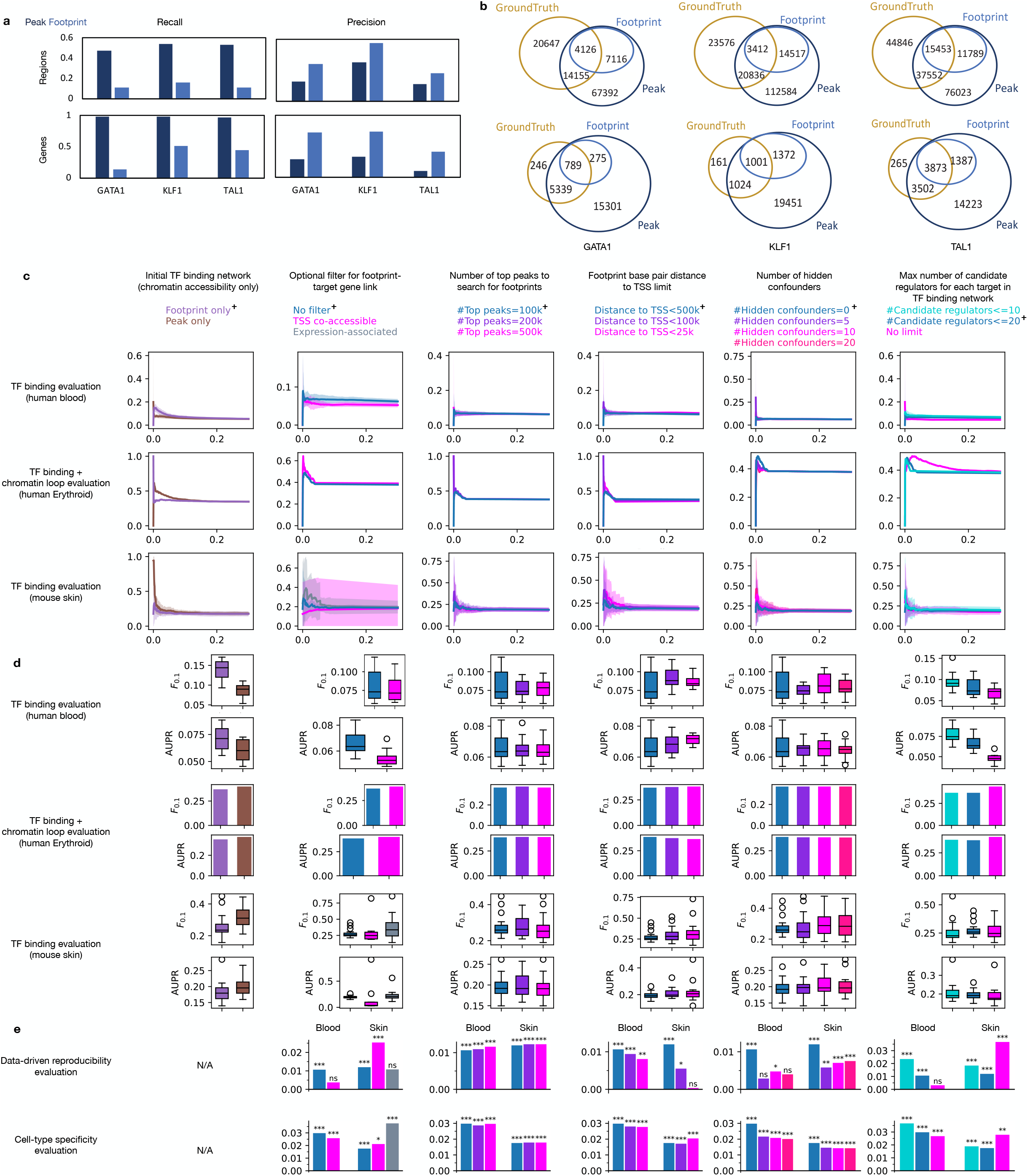
Benchmarking of different parameter choices within Dictys GRN reconstruction. **ab** Barplots of precision and recall (**a**) and Venn diagrams (**b**) of TF binding of chromosomal regions or their consequential annotation to nearby candidate target genes in the TF binding network for symbolic TFs in human Erythroid, when compared against the ChIP-seq + HiC results in Erythroid. **cde** TF binding and TF binding + chromatin loop evaluations’ Precision-Recall curves (**c**) and partial AUPR and F_0.1_ scores (**d**), and data-driven reproducibility and cell-type specificity evaluations’ (**e**) results for different Dictys settings and parameters. Cross annotation indicates the default parameter setting.

**Fig S5.**
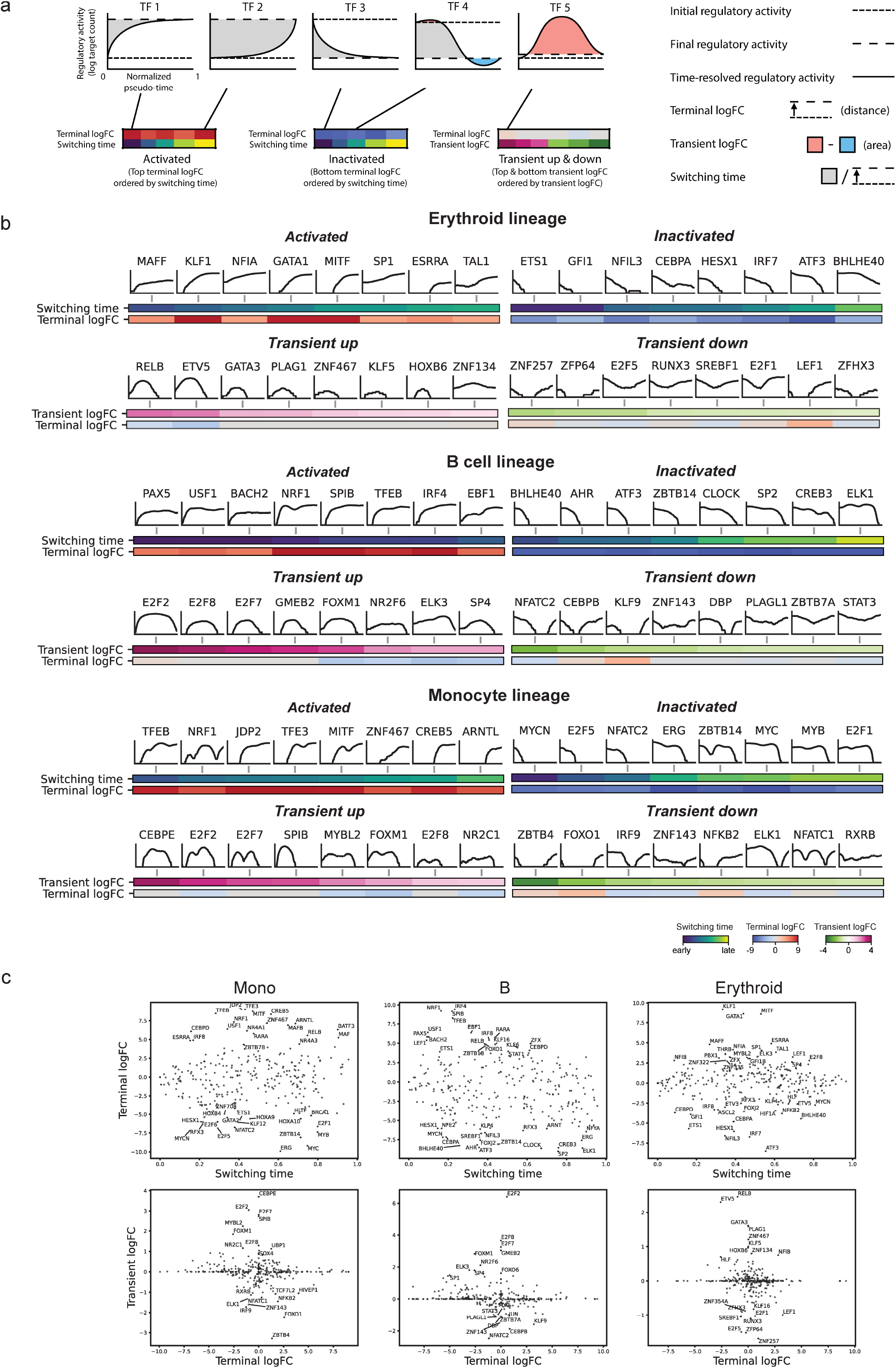
Dynamic GRNs from Dictys can be summarized into regulatory activity curve and its geometric features/characteristics to discover top TFs in four major patterns of regulatory activity shifts in human hematopoiesis. **a** Definition of three characteristic functions of regulatory activity curve from dynamic GRN and the unbiased TF discovery process for four patterns based on these characteristics. **b** Top 8 TFs detected for each regulatory activity pattern (Activated, Inactivated, Transient up, Transient down) from each lineage. **c** Scatter plots of these characteristics for all TFs across Erythroid, Monocyte and B cell lineages. Top TFs along Y axes are annotated.

**Fig S6.**
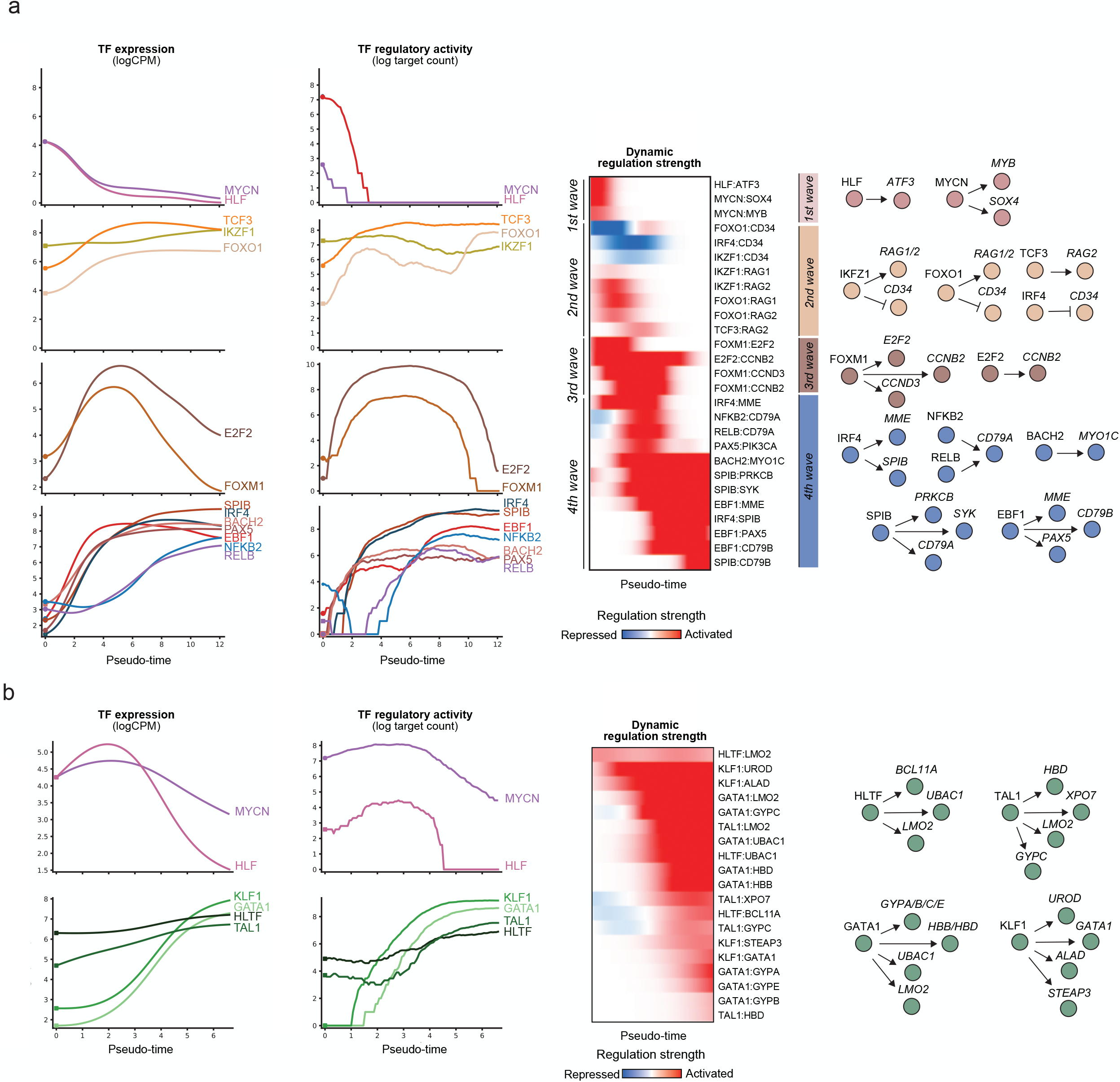
Dynamic GRNs identified established and novel TFs in human lymphopoiesis and erythropoiesis. Time-resolved dissection of master TFs and regulations during B cell (**a**) and Erythroid (**b**) development. Left: Expression and regulatory activity curves of TFs with established or novel involvement in the lineage. Center: Time-dependent strengths of individual regulations of lineage-defining markers by master TFs in left, as grouped in waves based on when each regulation peaks. Bold regulations are illustrated in right. Right: Schematic illustration of prominent regulations recovered in each wave.

Supplementary File 1: Extended texts, figures, and references

Supplementary Files 2-4: Movie visualization and analysis of dynamic GRNs for the Monocyte (2), Erythroid (3) and B cell (4) lineages. Top left: dynamic cell tracking highlighting those utilized for GRN reconstruction. Top right: dynamic differential analysis (as per **Fig. 2d**). Other panels (left to right): dynamic expression level, dynamic regulatory activity, dynamic regulation strength, dynamic GRN subnetwork (as per **Fig. 2f**). Rows: TFs coarsely grouped into different waves of regulation (one per row) across the developmental continuum.

Supplementary Table 1: Differential analysis logFCs and rankings

Supplementary Table 2: Subnetwork node and edge properties of presented TFs

Supplementary Table 3: Enrichr enrichment results

Supplementary Table 4: TF regulatory activity curves’ geometric characteristics for each lineage of human hematopoiesis

