## Supplementary File 1 for "Dictys: dynamic gene regulatory network dissects developmental continuum with single-cell multi-omics"

**3 Stochastic process network measured at steady state** **6**

### 1 Convention

In general, we use normal font for scalar variables, bold font for vectors, capitalized bold for matrices, and subscripts to indicate rows, columns, or elements of non-scalar variables. There are several exceptions:

- For conventionally capitalized variables, non-scalar forms are in capitalized bold font.
- For greek variables, neither bold font nor capitalization is applied regardless of dimensionality.

### 2 Brief overview

#### 2.1 Stochastic process networks

Consider $n$ variables measured over time $t$ as $\mathbf{x}(t) = (x_1(t), x_2(t), \ldots, x_n(t))^T$ where each $x$ corresponds to the true log expression level of each gene in gene regulatory network. Note here $\mathbf{x}(t)$ corresponds to their unknown true expression levels instead of their measured values. For the purpose of this paper, we consider $\mathbf{x}(t)$ to be log expression level so that in the linear regime, the model defined below would better capture the multiplicative nature of chemical master equations, although the model remains empirical.

Their interactions can be modeled at the time derivative (velocity for short) level with the stochastic differential equation (SDE) as[1]

$$\mathrm{d}x_i(t) = f_i(x_1(t), \ldots, x_n(t))\mathrm{d}t + \sum_j g_{ij}(x_1(t), \ldots, x_n(t))\mathrm{d}W_j(t), \tag{1}$$

where $W_j(t)$ is an independent Wiener process for each $j$. Its matrix form is

$$\mathrm{d}\mathbf{x}(t) = \mathbf{f}(\mathbf{x}(t))\mathrm{d}t + \mathbf{G}(\mathbf{x}(t))\mathrm{d}\mathbf{W}(t). \tag{2}$$

The first, deterministic *r.h.s* term encodes the stochastic process network that models the interactions between variables. The second, stochastic term models all sources of unknown perturbations, such as external noise, hidden variables, and stochasticity in chemical reaction.

We can define a network $\mathcal{G} \equiv (\mathcal{V}, \mathcal{E})$ to represent this system with a tuple of the set of vertices/nodes $\mathcal{V} = \{1, 2, \ldots, n\}$ and the set of edges $\mathcal{E} \subseteq \{i \to j \mid i, j = 1, \ldots, n\}$. Each node $i$ represents variable $x_i$ and has the time-dependent property based on its measurement $x_i(t)$. Each edge's presence defines the dependencies in Equation 2 as $i \to j \in \mathcal{E} \iff \frac{\partial f_j}{\partial x_i} \neq 0 \vee \exists k, \frac{\partial g_{jk}}{\partial x_i} \neq 0$. We also use $\mathbf{Pa}_i \equiv \{j \mid j \to i \in \mathcal{E}\}$ to indicate the set of parental/regulator nodes for node $i$.

For better understanding, consider a linear stochastic process network and constant contributions from stochasticity as an example, i.e.

$$\mathbf{f}(\mathbf{x}(t)) = \alpha + \beta^T \mathbf{x}(t), \tag{3}$$

$$\mathbf{G}(\mathbf{x}(t)) = \sigma \mathbf{I}_n. \tag{4}$$

Here $\alpha \in \mathbb{R}^n$ and $\beta \in \mathbb{R}^{n \times n}$ correspond to the intercept and network edge strengths respectively, and $\mathbf{I}_n$ is the identity matrix of size $n \times n$. Equation 2 then becomes

$$\mathrm{d}\mathbf{x}(t) = (\alpha + \beta^T \mathbf{x}(t))\mathrm{d}t + \sigma \mathrm{d}\mathbf{W}(t). \tag{5}$$

Therefore $\beta_{ij}$ is the effect of variable $i$ on variable $j$'s velocity. In this stochastic process network, the edge presence $i \to j \in \mathcal{E} \iff \beta_{ij} \neq 0$. Moreover, the value of $\beta_{ij}$ is a property of the edge $i \to j$.

#### 2.2 Steady-state Bayesian networks

An alternative scenario is that the same variables are measured only once but for multiple objects/samples as $\mathbf{X} = (x_{ij})$ for variables $i = 1, \ldots, n$ and samples $j = 1, \ldots, k$. One example is the destructive quantification of gene expression levels in single cells. Due to the lack of a concordant definition of (biological) time between samples, a popular study design is to acknowledge the kinetic and stochastic nature of the process but assume

[1] There are many studies with ordinary differential equations for a deterministic kinetic network. However, to prepare for the model Dictys uses, we derive the model for stochastic processes. Deterministic system is a special case of this derivation with $\mathbf{G}(\mathbf{x}(t)) = 0$.

the system already settles at its steady-state at every observation[2]. This allows to directly study and model the steady-state joint distribution $P(\mathbf{x})$ from observations $\mathbf{X}$ without specifying the kinetics in Equation 2. The steady-state assumption also agrees with many experimental designs such as differential gene expression between perturbed and unperturbed samples, where perturbed samples are given sufficient time to reach their perturbed homeostasis.

To model the dependencies between steady-state values of the variables, one popular approach is to similarly define a directed acyclic graph (DAG) $\mathcal{G} = (\mathcal{V}, \mathcal{E})$. Unlike stochastic process networks, edges here define variable dependencies through conditional distribution, as

$$P\left(x_i, \{x_j \mid j \in \mathbf{Pa}_i\}\right) = P\left(x_i \mid \{x_j \mid j \in \mathbf{Pa}_i\}\right) P\left(\{x_j \mid j \in \mathbf{Pa}_i\}\right). \tag{6}$$

Edges should indicate true dependencies, requiring

$$\forall k \in \mathbf{Pa}_i,\ P\left(x_i \mid \{x_j \mid j \in \mathbf{Pa}_i\}\right) \neq P\left(x_i \mid \{x_j \mid j \in \mathbf{Pa}_i \wedge j \neq k\}\right). \tag{7}$$

Because DAG does not contain any cycle, the joint distribution can be decomposed with the chain rule as

$$P(\mathbf{x}) = \prod_{i=1}^{n} P\left(x_i \mid \{x_j \mid j \in \mathbf{Pa}_i\}\right), \tag{8}$$

where for any independent variable $x_i$ with $\mathbf{Pa}_i = \emptyset$, its distribution is simply $P\left(x_i \mid \{x_j \mid j \in \mathbf{Pa}_i\}\right) = P(x_i)$.

Due to the generality here without specifying any distribution family or optimization goal, DAG-based steady-state network inference covers a wide range of scoring function and non-Gaussian conditional distributions. In this sense, many regression-based network inference models fall into this category, even implicitly.

The inference of such steady-state networks first needs to guarantee the satisfaction of DAG property. This requires a node ordering that could be estimated in advance [1] or fitted alongside other aspects of network inference. This is a challenge to steady-state network inference due to the tradeoff between computational complexity and accuracy. For simplicity, here we assume node ordering is given for steady-state networks. This challenge does not apply for stochastic process network which does not require the network to be a DAG.

Then, to reconstruct the steady-state network from the measurements $\mathbf{X}$, we need to find the optimal conditional distribution $P\left(x_i \mid \{x_j \mid j \in \mathbf{Pa}_i\}\right)$ from the given distribution family for each node/variable $i$, maximizing the given scoring function. Many scoring functions such as likelihood can be decomposed by Equation 8 into independent optimization subproblems for each node, with potential gains in algorithmic complexities in time and memory or convergence speed to the optimum.

For example, consider the steady-state network with edges $x_1 \to x_2 \to x_3$ and $x_1 \to x_3$. This decomposes the joint distribution into $P(x_1, x_2, x_3) = P(x_1)P(x_2 \mid x_1)P(x_3 \mid x_1, x_2)$. Let us further restrict conditional distributions to normal distribution whose mean linearly depends on other nodes, with

$$x_i \mid \{x_j \mid j \in \mathbf{Pa}_i\} \sim N(\alpha_i + \sum_j \beta_{ji} x_j, \sigma_i^2). \tag{9}$$

Using likelihood as the scoring function to maximize, the system reduces to three separate linear model problems

$$\begin{aligned} x_1 &= \alpha_1 + \varepsilon_1, \\ x_2 &= \alpha_2 + \beta_{12} x_1 + \varepsilon_2, \\ x_3 &= \alpha_3 + \beta_{13} x_1 + \beta_{23} x_2 + \varepsilon_3. \end{aligned}$$

Therefore, $\beta_{ij}$ encodes the steady-state effect node $i$ has on node $j$, as a property of edge $i \to j$.

a

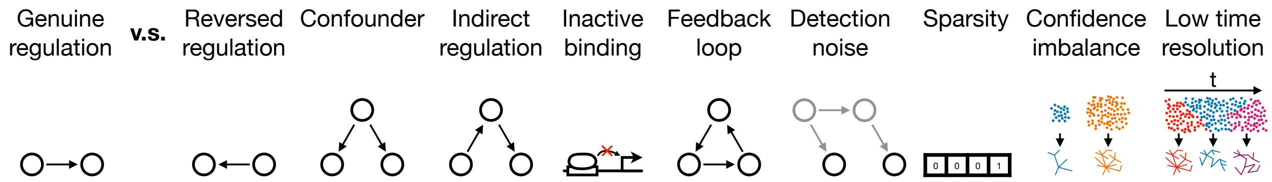

b

| | Origin | Challenge | Consequence | Our alleviation approach |
|---|---|---|---|---|
| Data | Transcriptome data | Reversed regulation<br>Confounder<br>Indirect regulation | False positive | Direct causal binding from epigenome |
| Data | Epigenome data | Inactive binding | False positive; no effect size | Regulatory effect from transcriptome |
| Model | Steady-state model | Feedback loop | Incompability | Stochastic kinetic network whose steady states are observed |
| Model | Regression model | Detection noise | Bias & variance in effect size; false negative | Probabilistic programming to model detection noise |
| Model | Single-cell data | Sparsity | Technical confounder; detection noise | Probabilistic modeling of scRNA-seq |
| Cluster | Sample imbalance | Confidence imbalance | Low comparability; sample size dependent variance | Fixed-size moving window in dynamic network |
| Cluster | Discrete cluster | Low time resolution | Missed key stages of network rewiring | Densely overlapped moving window along (pseudo-)time in dynamic network |

Figure 1: Challenges (**a**), their origin, consequences, and our alleviation approaches in Dictys (**b**).

#### 2.3 Challenges in steady-state network inference

Steady-state network inference has many challenges (Figure 1). They are typically due to the availability of data types[3], assumptions of the steady-state network model, preprocessing steps like clustering, or a combination of them. Below we give a brief overview in the context of gene regulatory network reconstruction and provide more details for challenges that are less mentioned in other network inference studies. Some other challenges also exist beyond the coverage of this paper, such as higher predictive powers in regression models do not necessarily indicate better knowledge recovery of causal relations.

##### 2.3.1 Reversed regulation

Even if all the model assumptions are valid and with infinite amount of data, Bayesian network may only be inferred up to the equivalence class of DAGs using steady-state samples of variables $x_i$ alone. Several different network structures may agree with the data best and equally well. Therefore, edge directions cannot be fully determined and genuine regulations cannot be fully distinguished from reversed regulations.

##### 2.3.2 Confounder

For the same reason, genuine regulations may be indistinguishable from the effects of a measured confounder. Unmeasured/unknown confounders add further challenges and can come from multiple sources such as imperfect assumptions of data.

[2]Non-steady-state Bayesian networks have been used to study variables at given time points. They have subtle differences in concept, causal structure (e.g. cycle-free), and statistical assumptions. They are not suitable for the studied question here due to the lack of time information.

[3]A simplistic formalization of biological terminologies in Figure 1 with network terms. Transcriptome data: samples of $x_i$. Epigenome data: network structure as a superset of edges $\tilde{\mathcal{E}} \supseteq \mathcal{E}$.

#### 2.3.3 Indirect regulation

Distinguishing indirect regulations through intermediate observed nodes from direct ones is challenging especially when the sample size is limited. This is even more challenging for unobserved intermediate nodes. A additional challenge is to utilize the existence of indirect regulations to improve network inference when the intermediate nodes are noisily measured.

#### 2.3.4 Inactive binding

Transcription Factors (TFs) regulate other genes' expressions by binding onto a nearby DNA region. However, binding is only a necessary condition for gene regulation. A large proportion of such bindings do not affect gene expression nearby, therefore here named inactive binding. Using such (epigenome) information alone can only yield a superset of allowed edges, but neither the true network nor the effect size (e.g. $\beta$) of the edges.

#### 2.3.5 Feedback loop

DAG-based steady-state network inference requires the network not to contain any cycles, as required by the chain rule of conditional distribution decomposition. For example, consider a two-node network with edges $\mathcal{E} = \{1 \to 2, 2 \to 1\}$. Obviously the chain rule (Equation 8) no longer holds:

$$P(x_1, x_2) \neq P(x_1 \mid x_2)P(x_2 \mid x_1). \tag{10}$$

The edge strength estimators would also become invalid and incur bias if still used.

Even when the true network does not contain any cycle, e.g. with $\mathcal{E} = \{1 \to 2\}$, negligence of the cycle-free property by naively assuming availability of all edges would also lead to the same incorrect model:

$$P(x_1, x_2) \neq P(x_1 \mid x_2)P(x_2 \mid x_1). \tag{11}$$

Therefore, DAG-based steady-state network inference cannot have any cycle and cannot model feedback loops in biology [2]. Although many methods can infer DAG-based steady-state networks while accounting for the cycle-free property [3, 4], many others methods simply ignored this challenge which would lead to biases in network inference. Moreover, feedback loops widely exist and have important roles in biology. Incompatibility with feedback loops will also introduce bias to the inferred network.

#### 2.3.6 Detection noise

Many biological measurements incur a detection noise. In quite some cases, detection noise is comparable with or even stronger than the original variable. ScRNA-seq is known to cause substantial measurement noises that bias the gene correlation and other analyses. Such detection noises can be modeled with an extra layer of variables $\mathbf{x}^{(\mathrm{obs})}$ dependent on the true value of expression level $\mathbf{x}$ with distribution $P(\mathbf{x}^{(\mathrm{obs})} \mid \mathbf{x})$. This is an extended version of steady-state network inference problem where edge existence is partially known/predetermined and some nodes are unobservable (without data).

However, most steady-state network inference models do not generalize to these two extensions easily. For example, regression models of network inference rarely account for detection noise. Ignoring detection noise is known to cause false positive and false negative edges in the inferred network [1].

#### 2.3.7 Sparsity

Single-cell RNA-seq datasets are known to be sparse. This introduces strong measurement noises that are cannot be modeled with Gaussian distribution; otherwise it introduces technical confounders [5]. ScRNA-seq sparsity also makes it hard to unbiasedly infer the variance of gene expression.

##### 2.3.8 Confidence imbalance

The amount of samples(/information) affects the variance of estimator of edge strengths, and consequently the confidence level of each edge. When inferring two networks separately for two groups of cells that are highly asymmetric in cell count, the imbalance in the confidence levels makes it challenging to compare the two networks. When using the same confidence threshold, two networks will have different edge sparsity levels. When using the same sparsity, the edges will have different confidence levels.

##### 2.3.9 Low time resolution

Clustering groups cells into non-overlapping subsets from each of which a network can be reconstructed. However, such coarse grouping provides a limited number of networks for typical questions that are insufficient for time-resolved processes such as cell differentiation, and may miss key stages such as turning or inflection points. These cell clusters also do not align perfectly along the differentiation path, which further reduces time resolution.

#### 2.4 Addressing these challenges

We address the reversed regulation, confounder, indirect regulation challenges by incorporating chromatin accessibility data, from which we can infer TF binding sites acting as a mechanistic indicator and necessary condition of direct causal effects (see main paper). We address the feedback loop challenge with stochastic process network which models the transcriptional rate (i.e. time derivative of variables) to allow for feedback loops as defined in Section 2.1, whose measurements are however performed at steady state (see Section 3). We account for detection noise and sparsity challenges with probabilistic modeling of scRNA-seq process (see Section 4). We handle sample imbalance and discrete cluster challenges with dynamic network that relies on a densely overlapped, fixed-size moving window to choose cell subset to reconstruct the network (see main paper).

### 3 Stochastic process network measured at steady state

We propose to address the indirect regulation and feedback loop challenges with a hybrid framework — a stochastic process network measured at steady state. The stochastic process network is defined in Equation 2, while its steady-state solution could be computed analytically. Because each variable only affects the time derivative of other variables, stochastic process network allows cycles or feedback loops. Although this model assumes the system to be at steady state when measured, it is identically assumed in the steady-state network. In this sense, this framework does not introduce any additional assumption. Instead, it no longer requires the network to be cycle-free, which is needed by steady-state Bayesian networks.

The stochastic process network can be mathematically formulated as infinite-sized chain graph models within the framework of non-steady-state Bayesian networks or structural equation models [6, 7]. However, the Bayesian network formulation does not provide network inference as for conventional Bayesian networks, suggesting that the hybrid framework is distinct from Bayesian networks.

Indirect effects are modeled by considering the expected stead-state expression changes of other genes when one gene's kinetic parameter (basal transcriptional rate) is perturbed. In this sense, we rely on the biological meaning of intervention in physical world to derive indirect effect size. It can automatically account for infinite propagations of variations on the network and utilize these indirect effects to optimize parameter(/network) inference. This is already described in the main paper.

We account for detection noise with probabilistic models of the technical detection process. For scRNA-seq, we model it as a binomial distribution from the steady-state distribution as previously found appropriate, as detailed in the main paper. The full generative process can be modeled with probabilistic programming and therefore the kinetic and stochastic parameters can be inferred from scRNA-seq read counts, as explicitly described in the next section. This provides a stochastic process network inference framework with demonstrated performance in high-dimensional complex systems with strong non-Gaussian measurement noise, among other challenges in Figure 1.

#### 3.1 Without regularization

For steady-state measurements of Equation 2, we replace time $t$ with hypothetical time $\tau$ without any predefined physical or biological meaning. At steady state, it approaches the $\tau \to \infty$ limit and therefore becomes absent in the final result. Consider the linear model without self-regulation as

$$\mathrm{d}\mathbf{x}(\tau) = \left(\alpha + \beta^T \mathbf{x}(\tau)\right) \mathrm{d}\tau + \sigma \mathrm{d}\mathbf{W}(\tau). \tag{12}$$

For gene regulatory network, $\alpha \in \mathbb{R}^n$ corresponds to basal transcriptional rate, and $\beta \in \mathbb{R}^{n\times n}$ with $\beta_{ii} = 0$ indicates the strength of individual edges of the network. The stochasticity coefficient can be decomposed into two submatrices

$$\sigma \equiv (\sigma^{(\mathrm{v})}, \sigma^{(\mathrm{cov})}). \tag{13}$$

The first submatrix includes per-gene stochasticity and unexplained variations as $\sigma^{(\mathrm{v})} \equiv diag(\sigma_1^{(\mathrm{v})}, \ldots, \sigma_n^{(\mathrm{v})})$. The second submatrix includes stochasticity and unexplained variations affecting multiple genes, here named hidden confounders, as $\sigma^{(\mathrm{cov})} \in \mathbb{R}^{n\times m}$, where $m$ is the number of hidden confounders.

Equation 12 is the SDE for multivariate Ornstein–Uhlenbeck (OU) process, whose steady-state solution can be analytically solved as

$$\mathbf{x}(\tau \to \infty) \sim MVN(\bar{\mathbf{x}}, \omega), \tag{14}$$

where $\bar{\mathbf{x}} \equiv -(\beta^T)^{-1}\alpha$ and $\omega$ is the solution of the continuous Lyapunov equation

$$\beta^T \omega + \omega \beta = -\sigma \sigma^T. \tag{15}$$

The solution for $\omega$ exists only when the real part of all $\beta$'s eigenvalues are negative. Otherwise, $\mathbf{x}(\tau)$ tends to diverge exponentially as $\tau \to \infty$.

#### 3.2 With regularization

The divergence at $\tau \to \infty$ represents positive feedback loops leading to modes that self-enhance exponentially over time. This causes other terms in Equation 12 to have diminishing relative contributions, including intercept and stochasticity. However, this divergence is neither biological nor physical, because each cell has limited production capacity for mRNAs and mass cannot grow infinitely in a limited space. To address that, we allow the intercept and stochasticity terms to match the divergence with an exponential scaling with time, as

$$\mathrm{d}\mathbf{x}(\tau) = \left(\alpha e^{\lambda\tau} + \beta^T \mathbf{x}(\tau)\right) \mathrm{d}\tau + e^{\lambda\tau} \sigma \mathrm{d}\mathbf{W}(\tau). \tag{16}$$

Switching variables to $\tilde{\mathbf{x}}(\tau) \equiv e^{-\lambda\tau}\mathbf{x}(\tau)$, we have

$$\mathrm{d}\left(e^{\lambda\tau}\tilde{\mathbf{x}}(\tau)\right) = \left(\alpha e^{\lambda\tau} + e^{\lambda\tau}\beta^T \tilde{\mathbf{x}}(\tau)\right) \mathrm{d}\tau + e^{\lambda\tau}\sigma \mathrm{d}\mathbf{W}(\tau), \tag{17}$$

i.e.

$$\mathrm{d}\tilde{\mathbf{x}}(\tau) = \left(\alpha + (\beta^T - \lambda\mathbf{I})\tilde{\mathbf{x}}(\tau)\right) \mathrm{d}\tau + \sigma \mathrm{d}\mathbf{W}(\tau), \tag{18}$$

where $\mathbf{I}$ is the identity matrix.

Renaming the variable $\tilde{\mathbf{x}}(\tau)$ to $\mathbf{x}(\tau)$, we get to a familiar form

$$\mathrm{d}\mathbf{x}(\tau) = \left(\alpha + (\beta^T - \lambda\mathbf{I})\mathbf{x}(\tau)\right) \mathrm{d}\tau + \sigma \mathrm{d}\mathbf{W}(\tau). \tag{19}$$

Therefore potential divergence due to positive eigenvalues in $\beta$ (i.e. positive feedback loops) can be offset with exponential time scaling. This regularization behaves effectively as a negative self-regulation in the network that, when large enough, can prevent divergence from positive eigenvalues.

Note that in Equation 19 no scale for time is specified by any variable. A time-scaling invariance exists which can be fixed by setting $\lambda = 1$ for simplicity and comparability between different networks (i.e. $\beta$) but without the loss of generality. Therefore, $\lambda = 1$ is set for this supplementary information. In Dictys, we set $\lambda$ as a free variable for easier parameter inference but nevertheless transform it to one with the time-scaling invariance before network analysis.

Because Equation 19 also fits in the OU process framework, we can perform similar derivations. Now

$$\mathbf{x}(\tau \to \infty) \sim MVN(\bar{\mathbf{x}}, \omega), \tag{20}$$

where $\bar{\mathbf{x}} \equiv \mathbf{B}^{-1}\alpha$, $\mathbf{B} \equiv \mathbf{I} - \beta^T$, and $\omega$ is the solution of the continuous Lyapunov equation

$$\mathbf{B}\omega + \omega\mathbf{B}^T = \sigma\sigma^T. \tag{21}$$

#### 3.3 Total effect steady-state network

The stochastic process network defines the direct effects between nodes. To account for effect propagation along the network edges, we can compute the total effect network that includes direct and indirect effects, by looking at the effect of perturbing basal transcriptional rate of one gene on other genes' mean expression at steady state. This is described in the main paper.

### 4 Probabilistic model for read count matrix

In this section we describe the probabilistic model for the scRNA-seq read count matrix, and link the variables in Dictys code to the variables in Section 3 where possible.

#### 4.1 With exact Lyapunov equation solver

The plate notation for the full probabilistic model is shown in Figure 2. For plates and dimensions:

- *edge* is all the edges allowed by the TF proximal binding network of potential gene regulations (Main Paper Figure 1**a**).
- *gene* is all genes in the cell subpopulation dataset after QC.
- *tf* is all TFs having at least one potential target in the TF bind network (Main Paper Figure 1**a**).
- *cell* is all cells in the subpopulation after QC.
- *pc0* is the low dimensions used to represent the offdiagonal terms in the covariance matrix of stochasticity to characterize hidden confounders.
- *pc* is the low dimensions used to represent the offdiagonal terms in the covariance matrix of biological gene variations (i.e. true gene expression). By default, it is the sum of the dimensions of pc0 and the number of TFs, so it can fully capture the covariance matrix without any loss of information.
- *confounder* is the list of known confounders whose valued are measured for each cell and whose effects on each gene will be estimated and accounted for.

For variables:

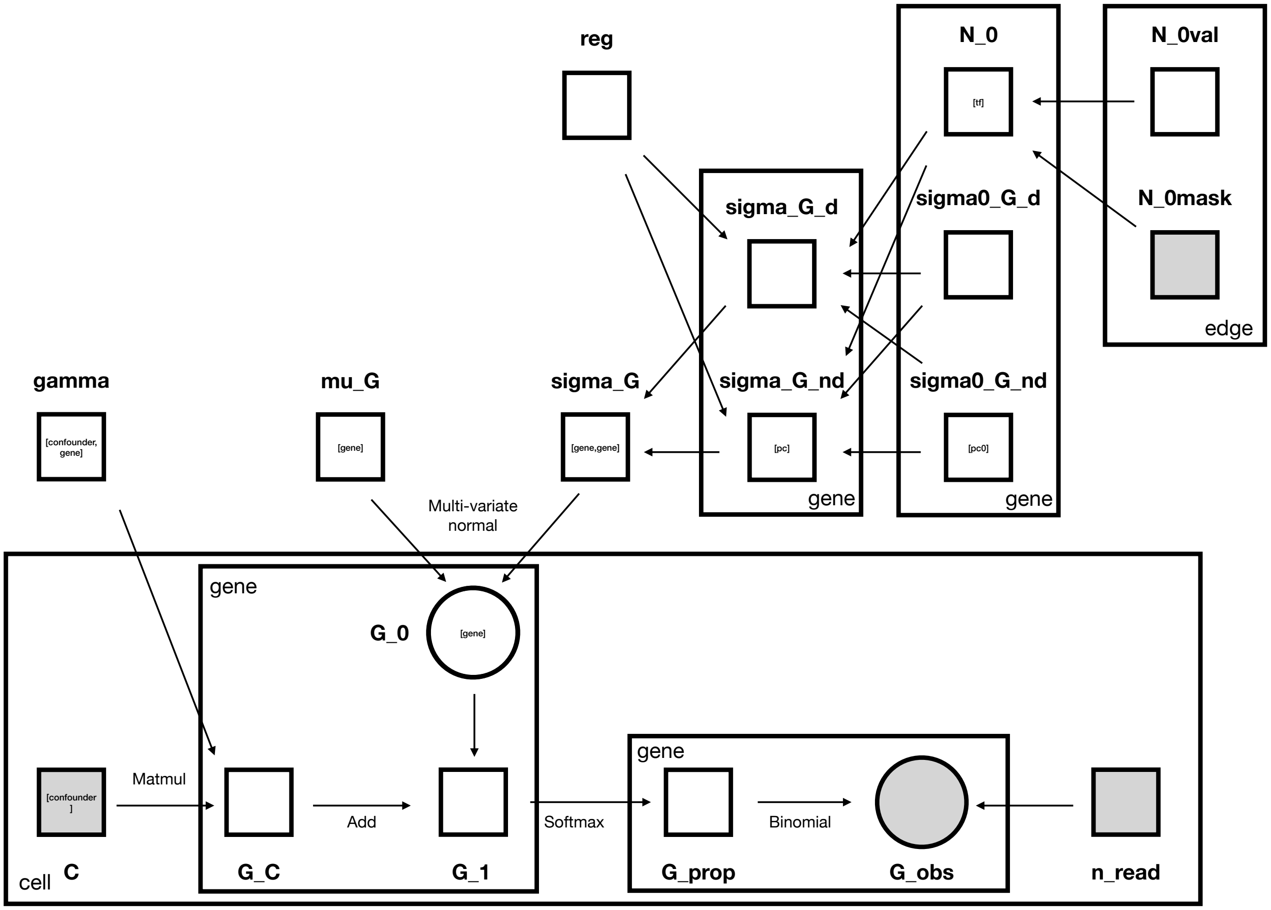

Figure 2: Plate notation of graphical model with an exact Lyapunov equation solver.

- *N_0mask* $\in \mathbb{N}_0^{n_{edge}\times 2}$ is the gene regulation edges allowed by the TF binding matrix, in terms of TF and gene indices. It is the index part of a sparse representation of the GRN matrix. Only these $n_{edge}$ edges can be present in the GRN matrix $\beta$.
- *N_0val* $\in \mathbb{R}^{n_{edge}}$ is strength of gene regulation for each corresponding edge in *N_0mask*. It is the value part of a sparse representation of the GRN matrix.
- *N_0* $\in \mathbb{R}^{n_{tf}\times n_{gene}}$ is the matrix form of GRN decompressed from the sparse representation with *N_0mask* and *N_0val*. It is a submatrix of $\beta$ as $\beta = (N\_0^T, \mathbf{0})^T$.
- *sigma0_G_d* $\in \mathbb{R}^{n_{gene}}$ is the strength of independent stochasticity applied on each gene. So $\sigma^{(\mathrm{v})} = diag(sigma0\_G\_d)$.
- *sigma0_G_nd* $= (\sigma^{(\mathrm{cov})})^T \in \mathbb{R}^{n_{pc0}\times n_{gene}}$ is the strength of inter-dependent stochasticity applied on each gene.
- *reg* $= \lambda \in \mathbb{R}^+$ is the regularization strength.
- *sigma_G_d* $\in \mathbb{R}^{n_{gene}}$ parameterizes the diagonal part of the covariance matrix of steady-state log gene expression.
- *sigma_G_nd* $\in \mathbb{R}^{n_{pc}\times n_{gene}}$ parameterizes the low-rank non-diagonal part of the covariance matrix of log steady-state gene expression.

- $sigma\_G = \omega = diag(sigma\_G\_d) + sigma\_G\_nd^T sigma\_G\_nd \in \mathbb{R}^{n_{gene} \times n_{gene}}$ is the full covariance matrix of steady-state log gene expression.

- $mu\_G = \bar{\mathbf{x}} \in \mathbb{R}^{n_{gene}}$ is the expectation of steady-state gene expression distribution.

- $G\_0 \in \mathbb{R}^{n_{gene} \times n_{cell}}$ is the unnormalized biological steady-state log expression level for each gene in each cell. For each column $i$, $G\_0_{\cdot,i} \sim i.i.d\ MVN(mu\_G, sigma\_G)$.

- $C \in \mathbb{R}^{n_{confounder} \times n_{cell}}$ is measured values of known confounder for each cell.

- $gamma \in \mathbb{R}^{n_{confounder} \times n_{gene}}$ is unknown linear effect size of each confounder on each gene's log expression.

- $G\_C = gamma^T C$ is aggregated confounder effect on gene expression.

- $G\_1 = G\_0 + G\_C$ is the unnormalized gene log expression after accounting for confounder effects.

- $G\_prop \in \mathbb{R}^{n_{gene} \times n_{cell}}$ is the normalized biological steady-state expression level for each gene in each cell. For each column $i$, $G\_prop_{j,i} = softmax_j(G\_1_{j,i})$.

- $n\_read \in \mathbb{N}_0^{n_{cell}}$ is the total number of reads in each cell, as obtained from data.

- $G\_obs \in \mathbb{N}_0^{n_{gene} \times n_{cell}}$ is the read count matrix for each gene in each cell, as in the data. $G\_obs_{i,j} \sim Binom(n\_read_j, G\_prop_{i,j})$.

The model contains hyperparameters and observations: *N_0mask, C, n_read, G_obs.* The model contains parameters to be inferred: *N_0val, sigma0_G_d, sigma0_G_nd, reg, mu_G, gamma.*

### 4.2 Invariances

The (loss function of the) full probabilistic model (Figure 2) is invariant under the constant translational transformation for all $i$:

$$\begin{aligned} x_i(\tau) &\to x_i(\tau) + \delta x, \\ \alpha_i &\to \alpha_i + \left(1 - \sum_{j \neq i} \beta_{ji}\right) \delta x. \end{aligned}$$

This translation leaves the SDE with $x_i(\tau) \to x_i(\tau) + \delta x$ on absolute abundance, but this constant translation is cancelled out in the scRNA-seq step as a binomial sampling process that is only dependent on their relative abundance. Therefore, the full probabilistic model is invariant under this translation, which is not a concern or interest in this study.

There are also transformations leaving the SDE (Equation 19) invariant but not the probabilistic model. Therefore, they do not affect this study so we just list several here.

- The SDE is invariant under the scaling transformation

$$\begin{aligned} x_i &\to \eta x_i, \\ \alpha_i &\to \eta \alpha_i, \\ \sigma_{ik} &\to \eta \sigma_{ik}. \end{aligned}$$

- The SDE is invariant under the scaling transformation for single gene $l$

$$\begin{aligned} x_l &\to \eta x_l, \\ \alpha_l &\to \eta \alpha_l, \\ \beta_{li} &\to \beta_{li}/\eta, \\ \sigma_{lk} &\to \eta \sigma_{lk}. \end{aligned}$$

#### 4.3 With approximate Lyapunov equation solver through additional loss

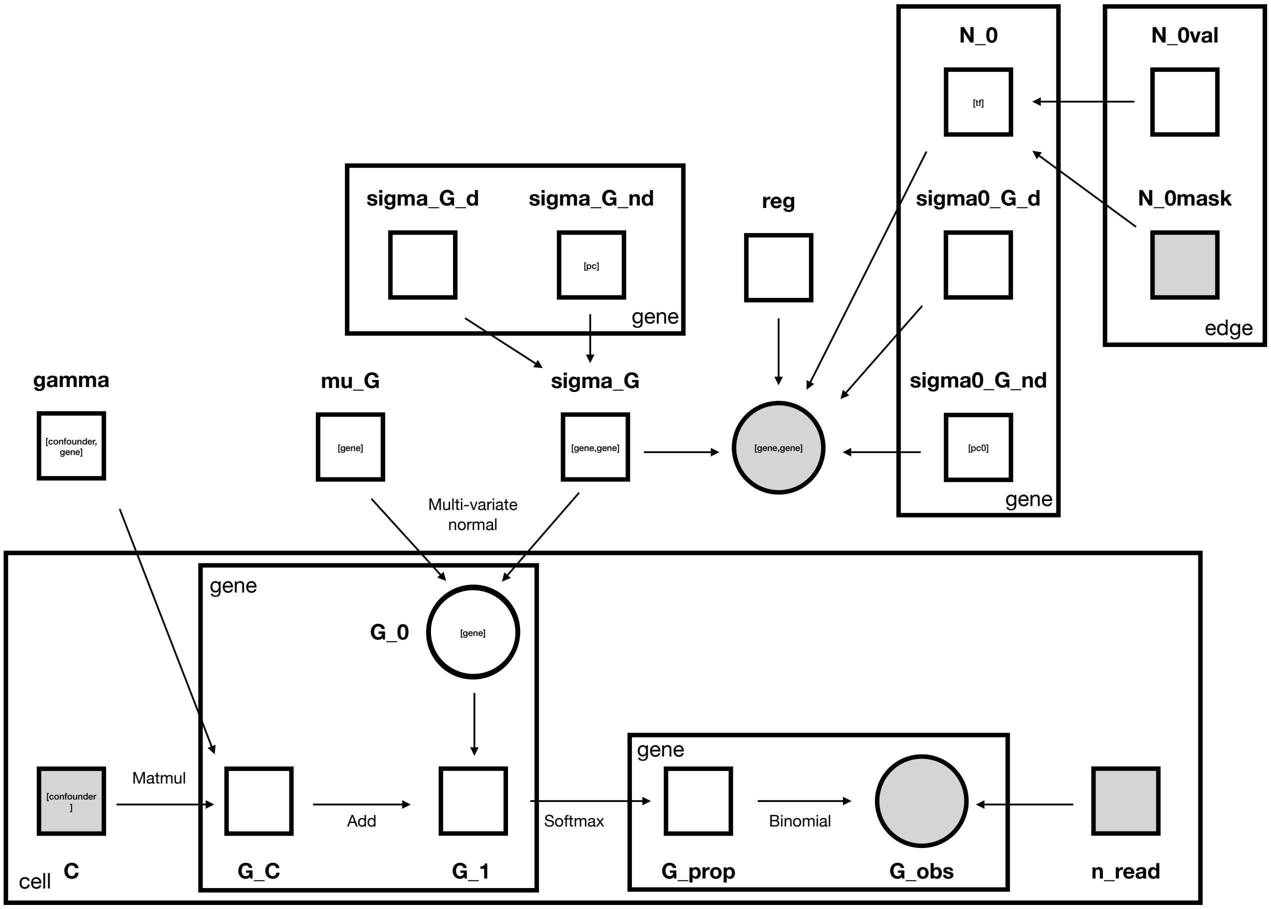

Figure 3: Plate notation of graphical model with an approximate Lyapunov equation solver through additional loss term.

However, a computational solver for Lyapunov equation is not publicly available on languages and platforms. It was infeasible to perform the same probabilistic programming on GPUs. To address this practical limitation, we introduced an extra loss term as in Figure 3.

The only difference arises in the variable *sigma_G_obs*$= \mathbf{B}\omega + \omega\mathbf{B}^T - \sigma\sigma^T$. When the Lyapunov equation solution is satisfied, we should have *sigma_G_obs*$=\mathbf{0}$. For a simple method to drive optimization towards this solution, we can construct an artificial observation for *sigma_G_obs* with a normal distribution of zero mean as

$$sigma\_G\_obs_{i,j} \sim i.i.d\ N(0, \sigma^2_{Lyapunov}), \tag{22}$$

where $\sigma_{Lyapunov}$ can be assigned a sufficiently large value for sufficient convergence to its solution. This is equivalent with a squared error loss on log likelihood.

#### 4.4 Additional considerations for approximate Lyapunov equation solver through additional loss

Replacing the Lyapunov equation solver with an extra loss term had two side effects which we addressed accordingly.

First, the change in variable dependency structure lost correct confidence level estimation. So we used the network edge strength $\beta$ for significance measure, instead of its confidence level, after post-processing scale-normalization to account for variance estimation bias.

Moreover, the extra loss term guarantees neither an exact solution for the stochastic process network nor consequently a finite solution for the steady-state network. To ensure convergence to a finite steady-state network, we included an adaptive post-inference regularization that further strengthens the regularization strength $\lambda$. Since the steady-state network computation requires matrix inversion $\mathbf{B}^{(\infty)} = \mathbf{B}^{-1}$ (Methods), we iteratively tested whether the largest eigenvalue of $\mathbf{B}$ stays below an upper bound (here 1.8). If not, a multiplier (here 1.1) is applied on $\lambda$ for stronger regularization, further reducing the diagonal elements of $\mathbf{B}$. This is performed iteratively until the largest eigenvalue of $\mathbf{B}$ stays in range.

### 5 Biology recovered from dynamic gene regulatory networks

#### 5.1 B cell lineage

Dynamic GRNs reconstruction was also performed across the B-cell trajectory in order to identify cell fate-determining TFs and their putative connections with prominent gene markers across B cell development (Suppl Fig6a). First, in agreement with the other two lineages, this analysis re-discovered the activity of pluripotency drivers at the progenitor states (HLF-ATF3, MYCN-MYB, MYCN-SOX4 [8, 9, 10, 11, 12]) to drop as part of the first wave. In the second wave, we found prominent lymphoid lineage regulators to supress the expression of the stem-cell marker CD34 (FOXO1-CD34, IKZF1-CD34, IRF4-CD34 [13, 14]), demarcating the initiation of cellular commitment towards B-cell development by exiting the progenitor (CD34+) states. At this same stage, we detected prominent activation of the RAG1/2 DNA recombinases, the main genes catalysing the VDJ recombination in pro-B/pre-B cells, by master TFs (IKZF1-RAG1/2, FOXO1-RAG1/2, TCF3-RAG2 [15]). Interestingly, we also detected subsequent regulations comprised of master cell cycle regulators activating cyclin genes (FOXM1-E2F2, E2F2-CCNB2, FOXM1-CCND3, FOXM1-CCNB2 [15]), demarcating the proliferative states found in pro-B and large pre-B cells. Finally, we detected a main regulatory "wave" comprised of master, B-cell development drivers which activate each other and prominent B-cell markers. These include IRF4-MME, IRF4-SPIB, NFKB2-CD79A, RELB-CD79A, PAX5-PIK3CA, BACH2-MYO1C, SPIB-SYK, SPIB-CD79B, EBF1-PAX5, EBF1-CD79B, EBF1-MME [16, 17, 18, 19, 20, 21]. Prominently, among these discovered connections, we identified previously established ones, including FOXO1-RAG1/2, TCF3-RAG2, E2F2-CCNB2, PAX5-PIK3CA, IKZF1-RAG1/2 and others [15, 19].

#### 5.2 Erythroid cell lineage

Following the trajectory towards erythroid development, we also identified the aforementioned regulators and regulatory connections underlying HSC and Progenitors programme (i.e. HLF, MYCN [8, 9]). In addition, we detected increased mRNA expression and regulatory activity of master TFs driving erythropoiesis, including GATA1, KLF1, TAL1 and HLTF [22, 23, 24, 25, 26] (Suppl Fig6b). Importantly, this analysis identified activation of genes defining the erythroid cell identity, including haemoglobin genes (HBB, HBD), genes involved in haemoglobin biosynthesis and metabolic processing (UROD, ALAD, STEAP3) [27], erythroid cell-defining surface markers (GYPA/B/C/E) [28], other prominent regulators (BCL11A, LMO2) [25, 24] and others (XPO7, UBAC1) [27]. As well as novel connections, this analysis confirmed the presence of previously known connections (i.e. KLF1-ALAD, KLF1-UROD, KLF1-STEAP3 [27], GATA1-HBB/HBD [29], GATA1-GYPB [30]).
